# Genome-Wide Association Study of Metabolic Traits in the Duckweed *Spirodela polyrhiza*

**DOI:** 10.1101/2024.07.26.605148

**Authors:** Martin Höfer, Martin Schäfer, Yangzi Wang, Samuel Wink, Shuqing Xu

## Abstract

The exceptionally high growth rate and high contents of flavonoids make the giant duckweed *Spirodela polyrhiza* (L.) Schleid. (Arales: Lemnaceae) an ideal organism for food production and metabolic engineering. To facilitate this, identification of the genetic basis underlying growth and metabolic traits is essential. Here, we characterized the genetics underpinning the growth and contents of 42 metabolites in *S. polyrhiza* using a genome-wide association (GWA) approach. We found that biomass positively correlates with contents of many free amino acids, including L-Glutamine, L-Tryptophane and L-Serine, but negatively correlates with many specialized metabolites, such as flavonoids. GWA analysis showed that several candidate genes were simultaneously associated with several metabolic traits, qualifying them as targets for growth manipulation and metabolic engineering. Together, this study provides insights into the metabolic diversity of *S. polyrhiza* and its underlying genetic architecture, paving the way for industrial applications of this plant via targeted breeding or genetic engineering.

## Introduction

With a biomass duplication rate of less than two days, *S. polyrhiza* is one of the world’s fastest-growing angiosperms (Ziegler et al. 2015). Due to its high levels of flavonoids and amino acids, *S. polyrhiza* is one of the best organisms for producing food resources and pharmaceuticals (Acosta et al. 2021; Baek et al. 2021; Zhao et al. 2015; Smith et al. 2024). To fully realise its industrial potential, specific improvements on its growth and metabolite production are needed, which requires a detailed understanding of the genetic mechanisms controlling these traits. Although previous forward genetic studies on metabolic traits in species like *Zea mays* L. (Poales: Poaceae) (Chen et al. 2016), *Oryza sativa* L. (Poales: Poaceae) (Chen et al. 2014; Zhang et al. 2019; Cu et al. 2021) and *Arabidopsis thaliana* (L.) Heynh. (Brassicales: Brassicaceae) (Angelovici et al. 2013; Angelovici et al. 2017) identified many promising candidate genes, the genetic principles controlling growth and plant metabolisms in *S. polyrhiza* largely remain unknown, hampering its further biotechnological optimization.

For most plants, to survive from natural stress factors, such as herbivory, they must carefully allocate their resources to growth or defense pathways. On a molecular level, free amino acids play a pivotal role in balancing resource distribution between biomass production and the synthesis of defense metabolites. Biosynthesis of amino acids can account for 50% of all carbon compound syntheses and consume 32% of total fixed carbon dioxide (Smith et al. 1961), making them a major carbon sink for biomass production in plants (Noctor and Foyer 1998). The central metabolic function of free amino acids is highlighted by their association with key metabolic pathways such as glycolysis, tri-carboxylic-acid (TCA) cycle, pentose-phosphate-pathway, urea-cycle and photorespiration (Figure 1) (Noctor and Foyer 1998). Besides their function in biomass gain, amino acids are precursors of many specialized metabolites like flavonoids and phenolic acids (Figure 1), which are involved in plant defense. Yet, the biosynthesis of specialized metabolites often negatively impacts plant biomass, primarily through the consumption of large nutrient resources (Züst et al. 2011) and autotoxicity (Dick et al. 2012; Li et al. 2021). Consequently, optimization for high crop yields often leads to high susceptibilities to pests due to a lack of defense metabolites, a phenomenon known as growth-defense trade-off (Züst and Agrawal 2017; Huot et al. 2014). On a regulatory level resource allocation from growth to defense pathways is controlled by phytohormones such as Jasmonic acid (JA), Abscisic acid (ABA), Salicylic acid (SA) and Auxin (IAA) (Huot et al. 2014; Hui et al. 2023; Aftab et al. 2010; Živanović et al. 2020a). Therefore, understanding the genetic mechanism controlling levels of phytohormones, free amino acids and related metabolites represents the key for optimizing the biomass yield and specialized metabolite contents in plants.

**Figure 1:**
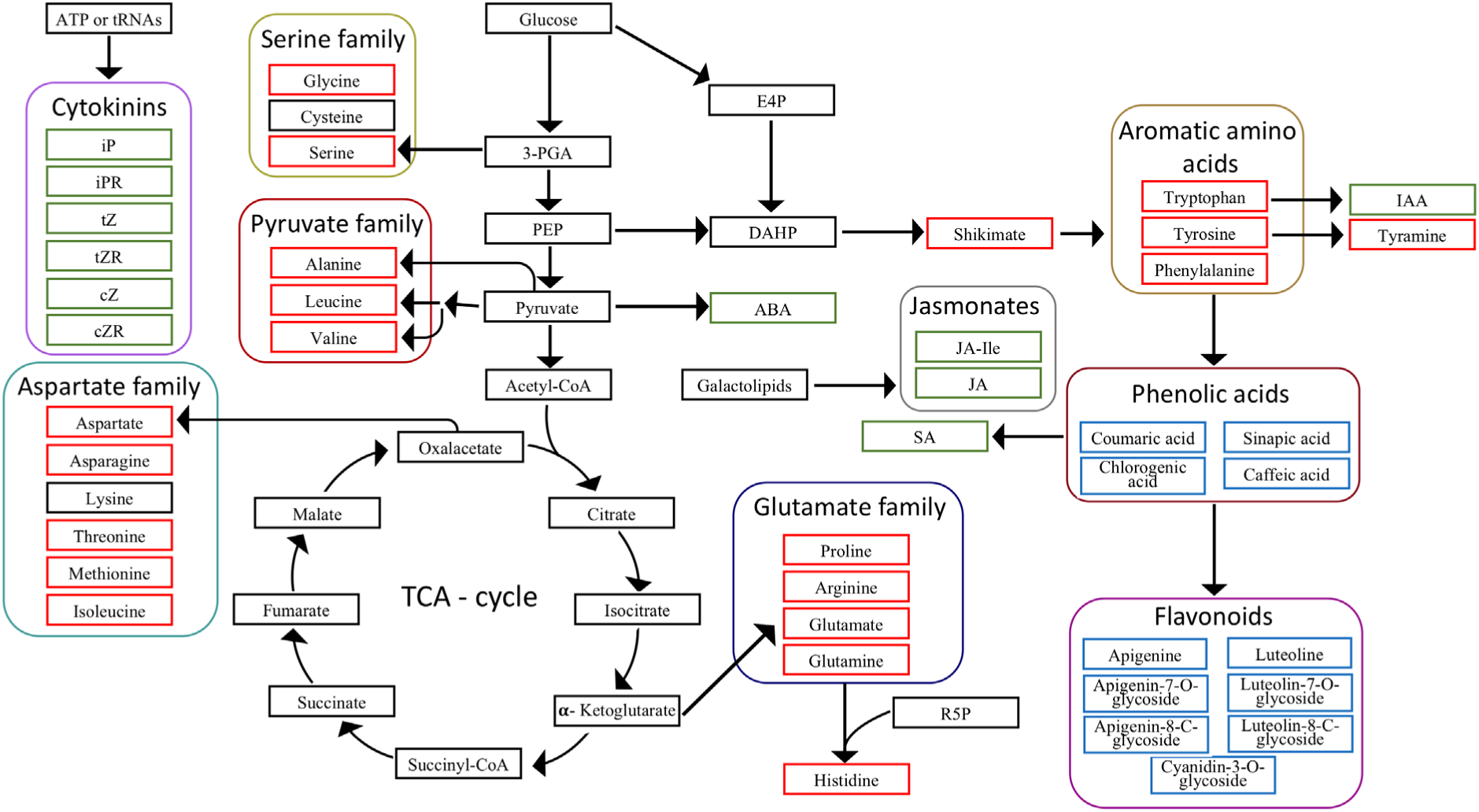
Overview of the biosynthesis of metabolites analyzed in this study. Metabolites were connected to their biochemical precursors with arrows. All quantified compounds were sorted into the three categories amino acid metabolism, specialized metabolism and phytohormones and shown in boxes colored red, blue, and green, respectively. Direct degradation products and biosynthetic precursors of amino acids such as tyramine and shikimic acid were grouped together with all quantified amino acids in the group amino acid metabolism. Amino acids were subcategorized into five families based on their last common precursor (serine, pyruvate, aspartate and glutamate) or on common chemical properties (aromatic amino acids). All specialized metabolites quantified in this study are phenylpropanoids derived from phenylalanine. Phenolic acids are also precursors of flavonoids. In contrast to amino acids and secondary metabolites, compounds with a major function in signalling were characterized as phytohormones. Chemically derived metabolites with the same precursors were further grouped in biosynthetic families, illustrated by frames of different colors. Abbreviations: TCA – tricarboxylic acid, iP – Isopentenyladenine, iPR – Isopentenyladenine riboside, tZ – trans-Zeatin, cZ – cis-Zeatin, tZR – trans-Zeatin riboside, cZR – cis-Zeatin riboside, 3-PGA – 3-Phosphoglycerate, PEP – Phosphoenolpyruvate, CoA - Coenzyme A, E4P – Erythrose-4-phosphate, R5P – Ribose-5-phosphate, DAHP – 3-Deoxyarabinoheptulonate-7-phosphate, JA – Jasmonic acid, JA-Ile – Jasmonic acid-Isoleucine conjugate, IAA – Indole-3-acetic acid, SA – Salicylic acid, ABA – Abscisic acid

Here, we focus on understanding the genetic control regulating levels of free amino acids and their related specialized metabolites in the giant duckweed. We quantified the contents of 42 metabolites (Figure 1) in 137 genotypes of *S. polyrhiza* and performed a genome-wide association study on the contents of these metabolites. We aim to address the following questions: 1) To what extent do growth and metabolism vary in *S. polyrhiza*? 2) which metabolites are associated with biomass and growth? and 3) which genes control these metabolic traits?

## Materials and Methods

### Data collection and sample preparation

In a previous study we estimated the growth of 138 *S. polyrhiza* genotypes under herbicide treatment and under control conditions (Höfer et al. 2024b). We quantified growth as relative growth rates (RGR) of frond area and frond number and as fresh weight and dry weight at the end of a seven-day growth period. For each genotype growth was estimated from three technical replicates. The quantified growth parameters are publicly available (Höfer et al. 2024a). A table documenting the geographic origins of all genotypes analyzed in this study can be found at: https://datadryad.org/stash/downloads/file_stream/2894056.

All analyses included in this study were based on plant material grown under control conditions (Höfer et al. 2024b). After harvesting, we freeze-dried plant material and stored it at room temperature (RT) till metabolite extraction. To have sufficient plant material for metabolite analysis, triplicates were pooled for each genotype. The RGRs were estimated (Höfer et al. 2024b) based on a published method (Ziegler et al. 2015), applying the formula: 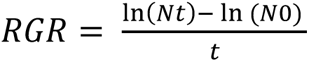, where *N_t_* is the frond number / area at the endpoint of the assay, *N_0_* is the frond number at the start of the cultivation and *t* as the duration of the bioassay.

### Metabolite extraction

To investigate genotypic variation in metabolite contents via LC-MS, we extracted 42 metabolites from pooled triplicates of 137 genotypes (Table S1, Figure 1). The number of analyzed genotypes was reduced from 138 to 137 due to mislabelling of one genotype, which was excluded from all analyses. The protocols used for LC-MS analysis of metabolite levels from different fractions are listed as Methods S1-5 in the supplemental part. The entire extraction procedure and quantification procedure is based on a method published previously (Schäfer et al. 2016).

The samples were ground with two steel balls at 30 Hz for 30 s in a TissueLyser (Qiagen, Venlo, Netherlands). Next, we aliquoted 10 mg of the homogenized samples in 96-well BioTubes (Axygen, New York, NY, USA). We then applied 800 µL of extraction buffer (methanol:water:formic acid (15:4:1) (v/v)) mixture containing [^13^C_6_]-IAA (1 ng), [^2^H_5_]-JA (10 ng), [^2^H_6_]-ABA (10 ng), [^2^H_4_]-SA (10 ng), [^2^H_5_]-tZ (0.5 ng), [^2^H_6_]-iP (0.2 ng) and [^2^H_6_]-iPR (0.2 ng) to each tube. We sealed the plate with a pre-cooled mat and homogenized the samples in a TissueLyser at 25 Hz for 60 s. Samples were incubated at -20 °C overnight. After that, we repeated the homogenization step at 25 Hz for 60 s. We then centrifuged the samples at 2250 g for 20 min at 4 °C and transferred 600 µL of the supernatant in new BioTubes, that were pre-cooled to -20 °C previously. The pellet was kept for re-extraction. We diluted an aliquot of 1 µL of the supernatant in 99 µL of algae amino acid standard mix -^13^C, ^15^N (0.5 ng/µL) (Sigma-Aldrich, St. Louis, MO, USA) and transferred the dilutions in 96-well PCR plates (Kisker, Steinfurt, Germany). The PCR plates were then stored at -20 °C till analysis of all amino acids and several secondary metabolites (Methods S1 and S2, Table S1). The pellet was reextracted with 600 µL of extraction buffer without standards. We homogenized the reextracted samples at 25 Hz for 60 s before incubation at -20 °C for 30 min. After another centrifugation step at 2250 g for 20 min at 4 °C, we mixed 600 µL of the supernatant from the reextracted samples with the remaining supernatant from the first part of the extraction. We centrifuged the combined supernatants at 2250 g for 20 min at 4 °C. Before loading our samples, we conditioned the HR-X columns (Macherey-Nagel, Düren, Germany) with 600 µL of MeOH, followed by 600 µL of extraction buffer and discarded the flow through. We collected the flow-through of our samples in Nunc 96-well Deep Well Plates (Thermo Fisher Scientific, Waltham, Massachusetts, USA). Next, we applied another 200 µL of extraction buffer and collected the flow through in the same well plate. We then evaporated the MeOH at 45 °C. Next, we added 850 µL of 1 N formic acid to each sample, mixed the samples at 20 Hz for 30 s using a TissueLyser and centrifuged them at 2250 g for 20 min at 4 °C. Before starting the next purification step, we conditioned HR-XC columns with 600 µL

MeOH and afterwards with 600 µL 1 N formic acid. The flow-through was discarded. We loaded the samples on the conditioned HR-XC columns and subsequently washed the columns with 1.2 mL of 1 N formic acid. We discarded the flow through and eluted the sample with 1 mL of 0.2 N formic acid in 80% (v/v) MeOH. We collected the eluates in BioTubes. After sample homogenization by inversion, we transferred 50 µL of the supernatant into a 96-well PCR plate for analysis of phytohormones (Method S3, Table S1). The sealed plates were stored at -20 °C till analysis.

For analysis of Indole-3-acetic acid (IAA) and phenolic acids (Method S4), the remaining eluate was evaporated under a constant nitrogen stream at 45 °C and samples were reconstituted to 50 µL with 0.2 N HCOOH in 80% (v/v) MeOH. We covered the plates before mixing at 20 Hz for 30 s. After centrifugation, samples were transferred to a 96-well PCR plate and were stored till analysis at -20 °C. For extraction of cytokinins (Method S5), we washed the HR-X column twice with 1 mL of 0.35 N NH_4_OH. Cytokinins were finally eluted with 1 mL of 0.35 N NH_4_OH in 60% (v/v) methanol. The samples were then dried under constant nitrogen flow. Next, we reconstituted sample volume to 50 µL with 0.1% (v/v) acetic acid. The samples were then homogenized at 20 Hz for 30 s, before incubation in the ultrasonic bath for 5 min. The samples were centrifuged at 2250 g for 20 min at 4 °C, and the supernatant was transferred to a 96-well PCR plate and stored at -20 °C until analysis (Method S5).

### LC-MS analysis of metabolite contents

We quantified all metabolites via LC-MS. For separation and quantification, we used a Nexera X3 UHPLC system (Shimadzu, Kyoto, Japan) coupled to an LCMS-8060 system (Shimadzu). For separation of all metabolites, we used a Zorbax RRHD Eclipse XDB-C_18_ column (50 x 3 mm, 1.8 µm) (Agilent, Santa Clara, CA, USA) with a 1290 Infinity II inline filter (0.3 µm) (Agilent). For all measurements, the autosampler was pre-cooled to 5 °C during the entire measurement procedure. As a mobile phase, we used a mixture of 0.05% (v/v) of formic acid and 0.1% (v/v) acetonitrile as solvent A and 100% methanol as solvent B, that were applied in gradient mode at a constant flow rate of 0.5 mL/min in all methods. The column oven temperature was always 42 °C. For analyses, we used an electrospray ionization source with the following parameters: nebulizing gas flow: 3 L/min, heating gas flow: 10 L/min, drying gas flow:10 L/min, interface temperature: 300 °C, DL temperature: 250 °C, heat block temperature: 400 °C and CID gas flow: 270 kPa. For metabolites measured in the positive and negative ionization modes, we applied interface voltages of 4000 V and - 3000 V, respectively (Table S2-6). A detailed description of individual metabolite analysis can be found in the supplemental (Method S1-5).

### Correlation and Principal Component Analysis

Next, we used regression and principal component analysis (PCA) to identify correlation patterns between metabolic parameters and to identify the influences of population structure on metabolite levels and growth. We established multiple correlations for all metabolite and growth data using the ggcorrplot R-package (version 0.1.4.1) (Kassambara 2023). We used the Pearson correlation coefficient to measure the strength of our linear correlations. All PCAs were conducted using the R package pcaMethods (version 1.88.0) (Stacklies et al. 2007). We calculated 95% confidence intervals based on standard deviations using the vegan package (version 2.6-2) (Oksanen et al. 2022) for all PCAs. All three-dimensional scatter plots were created using the R package scatterplot3d (version 0.3.42) (Ligges et al. 2003). For identifying metabolite categories, which strongly cluster together, we sorted all 42 metabolites into nine groups based on their chemical characteristics and biochemical precursors: Flavonoids, Phenolic acids, Aromatic amino acids, Glutamate family, Jasmonates, Serine family, Pyruvate family, Aspartate family and Cytokinins (Figure 1). The PCA on metabolite concentrations was subsequently done for all 137 genotypes.

Since *S. polyrhiza* has four genetic populations, as previously described by our lab (Wang et al. 2024; Xu et al. 2019), we next investigated to which extent differences in metabolite concentrations and growth are explained through population level. Assignment of the genotypes to different genetic populations was done in one of our previous studies (Wang et al. 2024). Due to the high proportion of clonal phenotypes in our accession, 137 genotypes were grouped into 97 clonal families for PCA on population structure of growth parameters and metabolite levels, respectively. For analysis, each clonal family was represented by the genotype with the highest sequencing coverage, as described previously (Höfer et al. 2024b).

### GWAS

To explore the genetic background underlying metabolic traits, we conducted GWAS on growth data and metabolite contents of all representative genotypes. We used single nucleotide polymorphisms (SNPs) and structural variations (SVs) (> 50 bp) as genetic markers for all GWAS analyses. To allow their usage for GWAS, SVs were recoded according to a previously published method (Lemay and Malle 2022). To correct for missing data, we performed an imputation of both SNP and SV data sets using beagle 5.4 version 22Jul22.46e (Browning et al. 2018).

We conducted GWAS on SNP and SV data using the vcf2gwas platform (version 0.8.7) (Vogt et al. 2021). Markers were pruned using the Plink software integrated into vcf2gwas, with phased r^2^ thresholds of 0.33 and 0.15 for SVs and SNPs, respectively. For filtering low abundant alleles, we applied a minor allele frequency (MAF) threshold of 5%, giving 10,057 SNPs and 1182 SVs for analysis. We corrected for population structure through principal component analysis (PCA) from the input genotype files. Here, we estimated population structure based on four PCs accounting for the four genetic populations of *S. polyrhiza*. All GWAS were conducted using a univariate linear mixed model (Zhou and Stephens 2012). The genotypic data, including the annotation, were published previously (Wang et al. 2024) and can be found under: https://github.com/Xu-lab-Evolution/Great_duckweed_popg (accessed on 22 March 2024).

### RT-qPCR quantification of candidate genes

Our GWAS detected a deletion in *SpUBP7* that was associated with increased contents of L-Glutamine and L-Serine. To validate the effect of the deletion on gene function, we studied the expression of *SpUBP7* (*SpGA2022_056000*) in two different allelic backgrounds.

For this we cultivated genotypes SP012 (low contents of L-Serine and L-Glutamine, no deletion in *SpUBP7*) and SP187 (high contents of L-Serine and L-Glutamine, homozygous for deletion in *SpUBP7*) in N-medium (Appenroth 2015) at 26 °C, 135 µmol photons · m^-2^ · s^-1^ 16 h/8 h light/dark rhythm. For each genotype, five replicates were made, each consisting of ten fronds growing as colonies. The fronds were cultivated in plastic beakers (Verpackungsbecher PP, transparent, round, 250 mL, Plastikbecher.de GmbH, Giengen, Germany), which were covered with perforated lids to allow gas exchange. Each beaker was filled with 150 mL N-medium.

After five days of cultivation, we harvested the root and frond tissue of each replicate separately for RNA extraction. We extracted RNA from < 20 mg fresh weight using the InnuPREP RNA mini kit (Analytik Jena, Jena, Germany). For each sample, RNA concentration was measured using a Nanodrop and integrity was checked via gel-electrophoresis on a 1% agarose gel. We used 600 ng of RNA for each cDNA synthesis following the instructions of the RevertAid First Strand cDNA synthesis kit (Takara, Shiga, Japan). For all cDNA syntheses, we used Oligo-dT primers. qPCRs were carried out using a RotorGene Q system (Qiagen, Venlo, The Netherlands) applying a time program with an initial denaturation of 98 °C – 3 min, followed by 40 cycles with 98 °C for 3 s and 60 °C for 20 s. Before qPCR, we estimated primer efficiency for *SpUBP7* by using serial dilutions of pooled cDNA templates from root and frond tissue samples. Primer specificity was checked by evaluating qPCR products on a 2% agarose gel and by melting curve analysis. We conducted all qPCR reactions according to the instructions of the KAPPA SYBR FAST kit (Roche, Basel, Switzerland). All cDNA samples were diluted in a 1:100 ratio in nuclease-free water before analysis. We used the alpha-elongation factor one gene *SpaEF* (*SpGA2022_005771*) and the glycerinaldehyd-3-phosphate dehydrogenase gene *SpGAPDH* (*SpGA2022_054082*) as references for visualization of candidate gene expression according to the delta-delta Ct method with multiple reference genes as published previously (Hellemans et al. 2007). The primers used for this qPCR study can be found in Table S7. The primers for the reference genes *SpaEF* and *SpGAPDH* were used in previous publications (Höfer et al. 2024b; Wang et al. 2024).

### Software and Statistics

We conducted all statistical analyses with R version 4.2.0. Standard errors of means were calculated using the plotrix R package (Lemon 2006). For all LC-MS measurements, we recorded and quantified analyte and standard peaks with LabSolutions software version 5.97. We estimated the broad sense heritability (H2.c) from our comprehensive data set, which includes repeatedly analyzed genotypes with the lme4 R package (Bates et al. 2015) following a method previously published by Cullis et al. (2006).

The significance of correlations was evaluated using the F-test. We used Bonferroni corrected *P*-value < 0.05 to determine significant genetic markers associated with our traits. The *P*-values for GWAS were calculated using the Wald test. In case of *SpUBP7*, we compared the SV effect of heterozygous and WT samples using a two-sided Student’s *t*-test. We used Students *t*-test for comparison of gene expression between two genotypes. We conducted the Levene test to check for homogeneity of variances using the car R package (Fox and Weisberg 2019).

## Results

### Intraspecific variation of growth rate and metabolic traits

We quantified 42 metabolites, including 20 free amino acids and their derivatives, 11 secondary metabolites and 11 phytohormones (Figure 1) among 137 *S. polyrhiza* genotypes. We found L-Asparagine and L-Glutamine were the most abundant free amino acid in *S. polyrhiza*, having average genotype tissue concentrations of 114.4 µmol/mg DW and 57.0 µmol/mg DW, respectively (Table S8). Abscisic acid (ABA) was the least abundant metabolite, showing an average concentration of 37 pmol/mg DW (Table S9).

Among all quantified metabolic features contents of flavonoids and phenolic acids showed the highest amounts of intraspecific variation (Table S10, Figure S1-4). Chlorogenic acid and Luteolin-7-O-glycoside showed 214.9 and 183.8-fold differences, respectively, among genotypes (Table S10, Figure S3). Growth rate, either quantified as RGRs of frond number or frond area, had less than 2.4-fold differences (Figure S1). On average, growth parameters also appeared to have a higher broad-sense heritability than metabolites (Table 1, Table S8-10). All growth parameters showed overall high broad-sense heritability values between 0.6 and 0.9, with fresh weight being the most heritable (Table 1). We found 25 metabolites reached high heritability values (H2.c > 0.3). The remaining 16 metabolites showed low heritability (H2.c < 0.3) (Table S8-10). Together, these findings suggest that growth rate and metabolic traits are variable within *S. polyrhiza*, and the majority of these traits are heritable.

**Table 1:** Phenotypic variation and heritability of fitness parameters in *S. polyrhiza*.

| Fitness trait | Mean | SE | Range |  | Fold change | H2.c |
| --- | --- | --- | --- | --- | --- | --- |
|  |  |  | Minimum | Maximum |  |  |
| Fresh weight | 227.7 mg | 7.3 mg | 71.6 mg | 507.7 mg | 10.4 | 0.865 |
| Dry weight | 16.67 mg | 0.5 mg | 3.4 mg | 35.2 mg | 10.4 | 0.602 |
| RGR of frond area | 0.323 d<sup>-1</sup> | 0.004 d<sup>-1</sup> | 0.165 d<sup>-1</sup> | 0.401 d<sup>-1</sup> | 2.4 | 0.841 |
| RGR of frond number | 0.259 d<sup>-1</sup> | 0.003 d<sup>-1</sup> | 0.152 d<sup>-1</sup> | 0.328 d<sup>-1</sup> | 2.2 | 0.802 |

Abbreviations: SE - standard error, H2.c – broad sense heritability according to Cullis et al. (2006)

### Metabolic traits correlate with biomass and growth

Plant metabolic traits are often associated with growth and biomass (Angelovici et al. 2013; Chen et al. 2016; Živanović et al. 2020b; Cu et al. 2021). In *S. polyrhiza*, we found plant metabolites showed higher correlations with biomass than with growth rate. In total, we found 19% and 29% of quantified metabolites showed significant correlation with growth rate (RGR of frond number and frond area) and biomass (fresh weight and dry weight) (Figure 2A, Figure S5 and S6), respectively. Among 20 quantified free amino acids, 11 showed a significant positive correlation with biomass (dry weight), including L-Glutamine, L-Tyrosine, L-Serine and L-Threonine (Figure 2A, Figure S6), whereas five were positively correlated with growth rate (RGR of frond number). Among phytohormones, only IAA and JA-Ile showed a positive correlation with biomass. Interestingly, most of the quantified specialized metabolites, while positively correlated with cytokinins, were negatively correlated with biomass (Figure 2A). Among all metabolites, Cyanidine-3-C-glycoside showed the strongest negative correlation with both growth rate and biomass (fresh weight) (Figure 2A, Figure S5 and S7).

**Figure 2:**
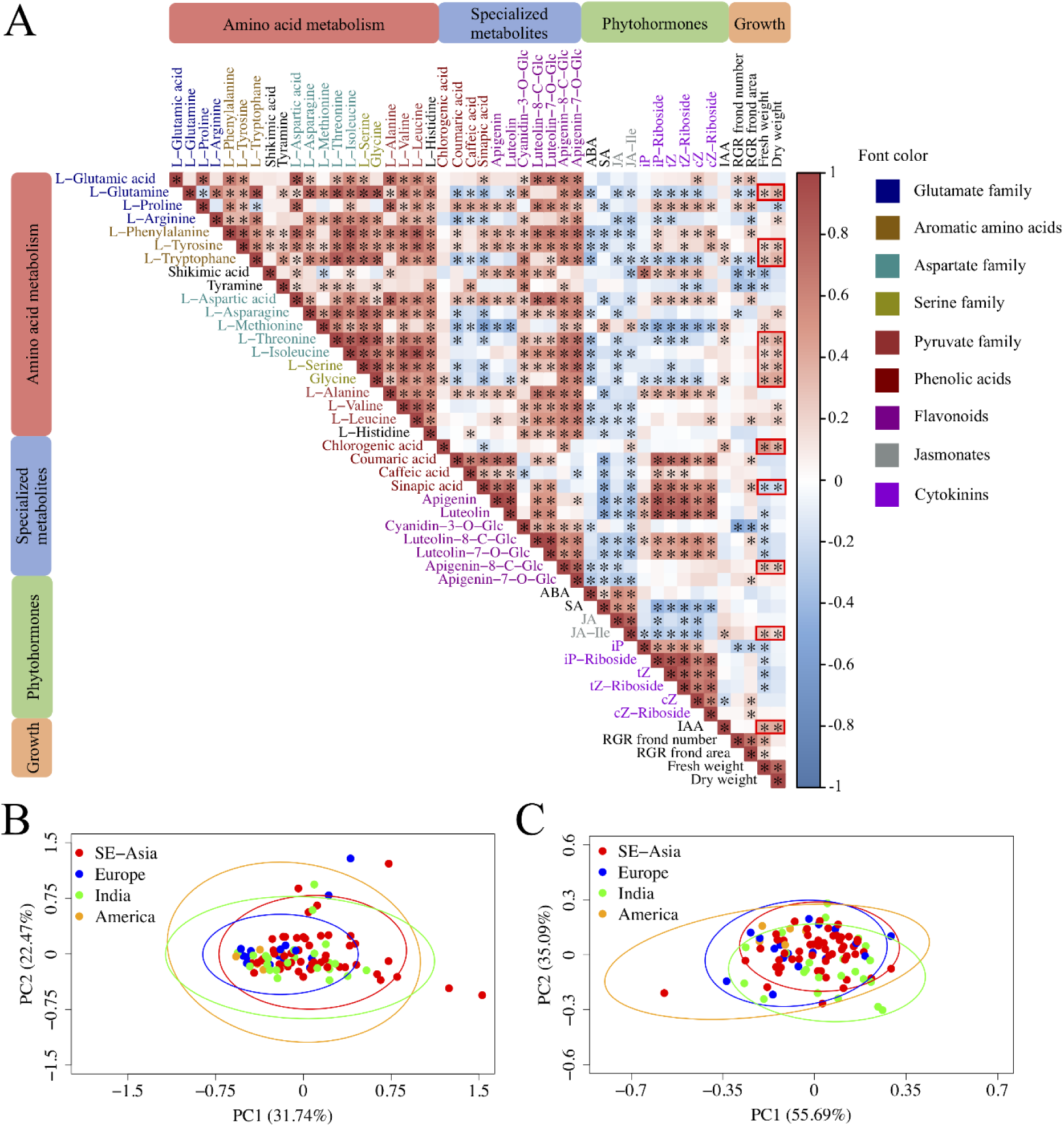
Correlation matrix (A) and principal coordinate analyses (PCA) on growth parameters (B) and metabolite content (C). A: Tissue concentrations of amino acids showed strong positive correlation patterns. In most cases, contents of amino acids were positively correlated with flavonoid content and biomass. Metabolite concentrations and growth rates were measured for 137 genotypes. All metabolites were categorized accordheing to Figure 1. The strength of the correlation was estimated using tPearson correlation coefficient and is reflected by the intensities of red color for positive and blue color for negative correlations. Asterisks highlight significant correlations (*F*-test, *P* < 0.05). Simultaneous correlations of dry and fresh weight were highlighted with red margins. B and C: population differences on growth (B) and metabolisms (C). All PCAs were conducted on 97 representative genotypes assigned to four previously defined genetic populations (Xu et al. 2019). The ellipses show 95 % confidence intervals calculated based on standard deviation. The color of the ellipses corresponds to the genetic populations. Abbreviations: iP – Isopentenyladenine, iPR – Isopentenyladenine riboside, tZ – trans-Zeatin, cZ – cis-Zeatin, tZR – trans-Zeatin riboside, cZR – cis-Zeatin riboside, JA – Jasmonic acid, JA-Ile – Jasmonic acid-Isoleucine conjugate, IAA – Indole-3-acetic acid, SA – Salicylic acid, ABA – Abscisic acid

Among the quantified metabolites, we found strong correlations within each metabolite group (Figure S8). Most of the free amino acids are positively correlated with each other and with flavonoids. Phenolic acids often were positively correlated with flavonoids (Figure 2A), reflecting their biosynthetic relationships (Figure 1).

Because *S. polyrhiza* has a strong population structure (Wang et al. 2024; Xu et al. 2019), which can confound the phenotypic variations and reduce the power of GWAS. We further investigated whether metabolisms differed among populations, using a principal component analysis (PCA). The results showed that there are no population-specific differences at the metabolic level (Figure 2B), indicating GWAS can be used for most of the metabolic traits in *S. polyrhiza*.

### GWAS of metabolite traits

We used a GWA approach to identify the genetic basis underlying different metabolic traits. Here, we focused on 10057 unlinked SNPs and 1182 SVs. A total of 75 SNPs and 15 SVs were significantly associated with the 42 metabolites (Table S11-13). Among them, 13 SNPs and three SVs were found to be associated with several metabolites, indicating their pleiotropic role in metabolisms. Surprisingly, we did not find any SNPs and only one SV, located in an intergenic region (> 30 Kb from next open reading frame), to be associated with growth rate and biomass (Table S12 and S13).

Among metabolic traits, we identified several loci that are associated with the contents of L-Glutamine, L-Tyrosine, L-Tryptophane, L-Serine, Chlorogenic acid and IAA. Most notably, several SNPs located within the *light harvesting protein 5* (*SpLHCB5*), which functions as a structural component of photosystem II (de Bianchi et al. 2008) were associated with contents of L-Glutamine, L-Valine, L-Tryptophan and L-Tyrosine (Figure 3 A-C, Table S13). Further, a SNP located upstream of a *U-box containing protein 4* (*SpPUB4*), which functions as a regulator of cell division processes in meristematic regions (Kinoshita et al. 2015), was associated with L-Tyrosine content. Three SNPs associated with the contents of L-Tyrosine, Chlorogenic acid and IAA were linked to *SpUGT89B1*, *SpCYP71AU50* and *SpMYBC1*, three genes functioning in secondary metabolite biosynthesis (Yamaguchi et al. 2014; Caputi et al. 2012; Ke et al. 2021) (Figure 3 B, D, E). A SNP within *SpDPE2*, which is functionally involved in starch metabolism is associated with IAA content (Li et al. 2022) (Figure 3E). For SVs, contents of L-Glutamine and L-Serine were associated with a 94 bp intronic deletion in the *Ubiquitin-carboxyterminal hydrolase 7 SpUBP7* (Figure 4A-C). Homologs of *SpUBP7* were shown to stabilize ubiquitin upon proteasome binding and regulate proteasomal activity (Leggett et al. 2002; Wu et al. 2019).The presence of the intronic deletion was associated with increased contents of L-Glutamine and L-Serine (Figure 4D and E) and an increased expression of *SpUBP7* in roots (Figure 4F and G). These findings suggest that biomass and metabolite contents are coordinated jointly through photosynthesis, starch metabolism, cell division, secondary metabolite biosynthesis and protein degradation.

**Figure 3:**
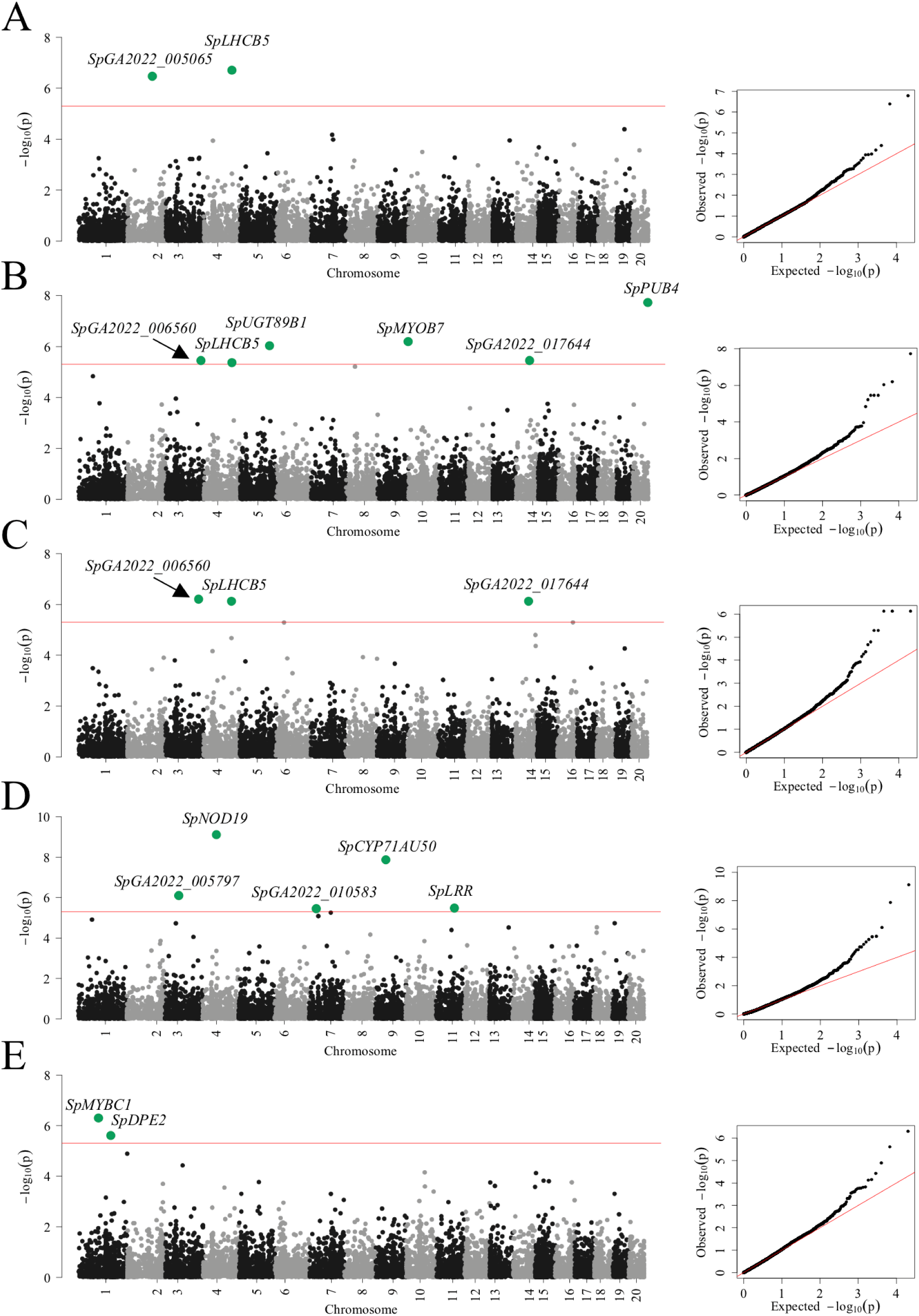
Manhattan plots of SNP-based GWAS for free amino acids. Each plot shows the GWAS result on the concentrations of L-Glutamine (A), L-Tyrosine (B), L-Tryptophan (C) and Chlorogenic acid (D) and Indole-3-acetic acid (IAA) (E) with their corresponding QQ-plots on the right side. The Bonferroni corrected *P*-value = 0.05 (Wald-test) at 4.97 · 10^-6^ was used as significance threshold and is shown as red line. All significant markers are labelled with the names of gene candidates.

**Figure 4:**
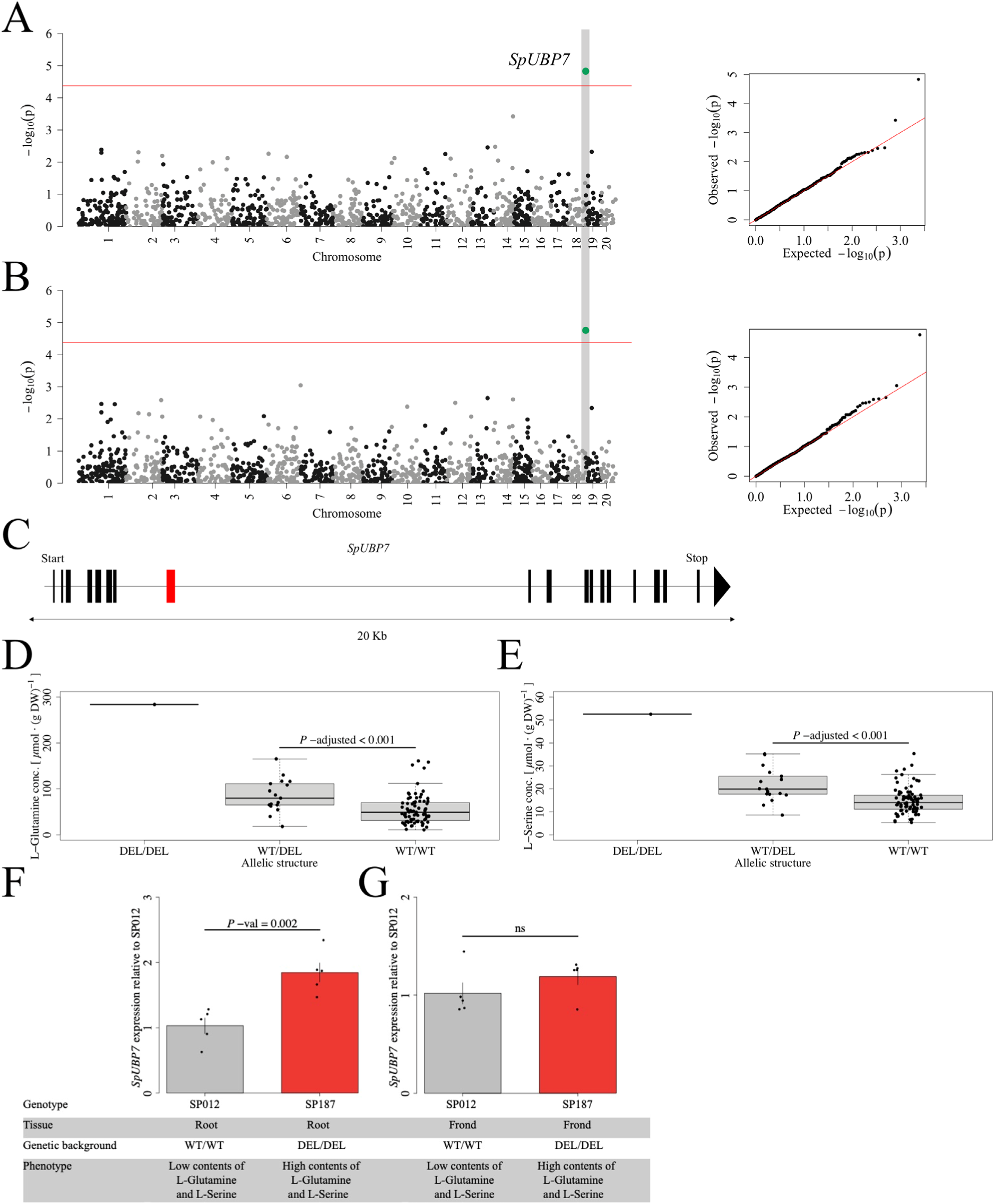
The presence of an intronic deletion in *SpUBP7* is associated with increased contents of L-Glutamine and L-Serine. A and B: The SV-based GWAS on L-Glutamine (A) and L-Serine (B). The red line represents the Bonferroni corrected *P*-value = 0.05 (Wald-test) at 4.23 · 10^-5^. C: The associated 94 bp deletion (red bar) is located within the 7^th^ intron of *SpUBP7*. D and E: The presence of the deletion was associated with increased contents in L-Glutamine (D) and L-Serine (E). F: *SpUBP7* shows genotype-dependent expression differences on root level. SP187, a genotype that is homozygous for the deletion, shows enhanced *SpUBP7* expression compared to genotype SP012 that lacks the deletion. G: No significant differential expression was observed between SP187 and SP012 in the fronds.

## Discussion

Using targeted metabolomics on 137 *S. polyrhiza* genotypes, we identified strong positive correlations between chemically related metabolites. Additionally, candidate genes involved in photosynthesis (*SpLHCB5*), protein degradation (*SpUBP7*) and organ development (*SpLOB*, *SpAGL62*) were repeatedly associated with different metabolite levels, suggesting strong genetic co-regulation. Since levels of free amino acids and secondary metabolites correlated with plant biomass in many cases, some of these candidate genes are of potential use for joint optimization of plant growth and nutrient level.

Free amino acids have large pool sizes and turnover rates in duckweed and were shown to serve as major carbon and nitrogen sources for starch and storage protein production (Evans et al. 2018). Previous work showed that supplementation of growth medium with certain amino acids like L-Glutamine was associated with increased protein content and biomass in duckweed (Shi et al. 2023). In agreement with these observations, the associations between free amino acid levels and biomass are mostly positive. Compared to free amino acids, phenylpropanoids and phytohormones are characterized by a higher functional diversity. In *S. polyrhiza*, anthocyanins were shown to alleviate Cr(VI) stress (Oláh et al. 2009), whereas luteolins and apigenins are involved in stress responses to copper and UV-light, respectively (Böttner et al. 2021). Previous studies found the contents of these three flavonoid classes to be inversely correlated with biomass in *S. polyrhiza* (Böttner et al. 2021; Oláh et al. 2009) and explained this by increased allocation of carbon resources to flavonoid biosynthesis (Oláh et al. 2009). Compared to this, our study revealed a more divers association pattern of flavonoid levels with biomass. While the contents of Sinapic acid, luteolins and Cyanidin-3O-glycoside were negatively associated with biomass, contents of Apigenin-8-C-glycoside and Chlorogenic acid showed positive correlations with biomass. In other plant species, changes in profiles of glycosylated flavonoids were shown to regulate plant growth by influencing auxin transport (Ringli et al. 2008), possibly explaining different association patterns of glycosylated luteolins and apigenins to plant biomass. Contents of chlorogenic acid were associated with increased leaf growth and increased contents of antioxidants and proteins, suggesting a regulatory function in plant growth (Zhang et al. 2024). The overall positive associations of cytokinin levels with most phenylpropanoids, were previously explained by their function in coordinating resource allocation to defense metabolism in plant development (Brütting et al. 2017). For phytohormones, levels of JA-Ile were positively correlated with biomass. In duckweeds, jasmonates show bimodal dose-response patterns, with small concentrations stimulating growth and turion germination and high concentrations delaying growth (Appenroth et al. 1991; Piotrowska et al. 2010). Since metabolites were measured in unstressed plant material, JA contents likely never exceeded constitutive levels, which would be required for significant growth inhibition. Together these correlation patterns suggest that *S. polyrhiza* can be simultaneously optimized for higher biomass and increased contents of certain phenylpropanoids.

On a genetic level, our study identified genetic loci that are associated with most secondary metabolite and phytohormone contents, whereas for many free amino acids, no significant associations were found. For growth rate and biomass, we did not find significant associations for any markers located within the genic region, likely due to their highly polygenic feature. Previous GWAS conducted on complex traits highlighted, that the success in identifying genetic associations greatly depends on sample size and phenotypic variation (Schizophrenia Working Group of the Psychatric Genomics Consortium 2014; Duncan et al. 2019; Chen et al. 2014). In line with previous reports (Chen et al. 2016; Pham et al. 2023), we found that secondary metabolite levels showed the highest average phenotypic variation across all metabolic traits, whereas variation for growth parameters was lowest. For secondary metabolites, high variation in tissue concentration was explained by their low trait complexity (Pham et al. 2023; Haghi et al. 2022; Chen et al. 2014). Whereas primary metabolites and growth seem to be regulated by many small effect loci, secondary metabolites are often controlled by just a few major-effect loci (Chen et al. 2014; Chen et al. 2016). Since small effect loci are often diffusely distributed across the genome, GWAS on primary metabolite levels and growth often require large sample sizes to detect them (Duncan et al. 2019). Compared to other association studies on plant metabolic traits (Chen et al. 2014; Chen et al. 2016; Cu et al. 2021; Angelovici et al. 2013), our GWAS suffered from a low sample size, likely explaining the lack of genetic associations for growth traits and several metabolite contents.

On a physiological level, differences in growth were correlated with the photosynthesis rate for duckweed (Sree et al. 2015). Connected to this, the metabolism of storage molecules plays a major role in plant growth. Interestingly, many biomass-correlated metabolites were associated with genes influencing photosynthetic efficiencies (*SpLHCB6*), protein turn over (*SpUBP7*), starch metabolism (*SpDPE*) or cell-cycle control (*SpPUB4*). These genes provide suitable candidates for future reverse genetic studies, that will be required for further elucidation of their exact function in duckweed metabolism. Taken together, this study provides insights into the molecular basis of growth regulation and metabolite homeostasis in *S. polyrhiza*. Through the identification of gene candidates associated with metabolic traits, this study contributes to laying the foundations for further optimization strategies in *S. polyrhiza*.

## Acknowledgement

We thank Marie Serwaty-Sárazová for maintaining the giant duckweed collection. This project was funded by the German Research Foundation (project numbers 427577435 to SX). The LC-MS/MS instrument was funded by the German Research Foundation (project number 435681637 to SX).

## Data availability statement

All raw data and scripts used for analysis can be found under: https://doi.org/10.17632/xwsxpfcysd.1 or https://data.mendeley.com/preview/xwsxpfcysd?a=2d1b7639-12ef-4665-bc72-5096a149004e

## Author contributions

MH analyzed the data. MH, MS, SW and YW performed the experiments. YW and SX provided resources and technical infrastructure. SX and MH conceived and supervised the project. MH wrote the manuscript. All authors contributed to the final version of the manuscript.

