## supplemental tables for "Genome-Wide Association Study of Metabolic Traits in the Duckweed *Spirodela polyrhiza*"

### Supplementary tables

Table S1: List of all measured metabolites with their metabolic classification. The numbers listed in the Method column correspond to the quantification protocols listed in the Supplemental methods part.

| Metabolite | Class | Method |
|---|---|---|
| **L-Alanine** | Amino acid metabolism | 1 |
| **L-Arginine** | Amino acid metabolism | 1 |
| **L-Asparagine** | Amino acid metabolism | 1 |
| **L-Aspartic acid** | Amino acid metabolism | 1 |
| **L-Glutamic acid** | Amino acid metabolism | 1 |
| **L-Glutamine** | Amino acid metabolism | 1 |
| **L-Isoleucine** | Amino acid metabolism | 1 |
| **L-Leucine** | Amino acid metabolism | 1 |
| **L-Phenylalanine** | Amino acid metabolism | 1 |
| **L-Proline** | Amino acid metabolism | 1 |
| **L-Serine** | Amino acid metabolism | 1 |
| **L-Threonine** | Amino acid metabolism | 1 |
| **L-Tryptophan** | Amino acid metabolism | 1 |
| **L-Valine** | Amino acid metabolism | 1 |
| **Chlorogenic acid** | Specialized metabolites | 1 |
| **Cyanidin-3-O-glycoside** | Specialized metabolites | 1 |
| **Luteolin-8-C-glycoside** | Specialized metabolites | 1 |
| **Luteolin-7-O-glycoside** | Specialized metabolites | 1 |
| **Apigenin-8-C-glycoside** | Specialized metabolites | 1 |
| **Apigenin-7-O-glycoside** | Specialized metabolites | 1 |
| **Glycine** | Amino acid metabolism | 2 |
| **L-Histidine** | Amino acid metabolism | 2 |
| **L-Methionine** | Amino acid metabolism | 2 |
| **L-Tyrosine** | Amino acid metabolism | 2 |
| **Tyramine** | Amino acid metabolism | 2 |
| **Shikimic acid** | Amino acid metabolism | 2 |
| **Apigenin** | Specialized metabolites | 2 |
| **Luteolin** | Specialized metabolites | 2 |
| **ABA** | Phytohormones | 3 |
| **SA** | Phytohormones | 3 |

| | | |
|---|---|---|
| **JA** | Phytohormones | 3 |
| **JA-Ile** | Phytohormones | 3 |
| **Coumaric acid** | Specialized metabolites | 4 |
| **Caffeic acid** | Specialized metabolites | 4 |
| **Sinapic acid** | Specialized metabolites | 4 |
| **IAA** | Phytohormones | 4 |
| **iP** | Phytohormones | 5 |
| **iPR** | Phytohormones | 5 |
| **tZ** | Phytohormones | 5 |
| **tZR** | Phytohormones | 5 |
| **cZ** | Phytohormones | 5 |
| **cZR** | Phytohormones | 5 |

Abbreviations: ABA – Abscisic acid, cZ – cis-Zeatin, cZR – cis-Zeatin riboside, IAA – Indoleacetic acid, iP – Isopentenyladenine, iPR – Isopentenyladenine riboside, JA – Jasmonic acid, JA-Ile – Jasmonic acid-isoleucine conjugate, SA – Salicylic acid, tZ – trans-Zeatin, tZR – trans-Zeatin riboside

Table S2: Technical parameters used for quantification of the amino acids and secondary metabolites quantified with Method S1 via LC-MS.

| Metabolite | Ionization mode | Start [min] | End [min] | Q1 [m/z] | Q3 [m/z] | Dwell time [ms] | CE | Q1/Q3 Pre Bias [V] |
|---|---|---|---|---|---|---|---|---|
| **L-Alanine** | + | 0.20 | 0.85 | 90.05 | 44.20 | 20 | -13 | -10/-18 |
| **L-Arginine** | + | 0.20 | 0.85 | 175.12 | 60.20 | 20 | -14 | -20/-24 |
| **L-Aspartate** | + | 0.20 | 0.85 | 134.04 | 88.20 | 20 | -12 | -13/-16 |
| **L-Glutamate** | + | 0.20 | 0.85 | 148.06 | 102.15 | 20 | -13 | -10/-19 |
| **[$^{13}C_3$, $^{15}N_1$]-Alanine** | + | 0.20 | 0.85 | 94.06 | 47.20 | 20 | -13 | -10/-18 |
| **[$^{13}C_6$, $^{15}N_4$]-Arginine** | + | 0.20 | 0.85 | 185.13 | 64.15 | 20 | -14 | -20/-24 |
| **[$^{13}C_4$, $^{15}N_n$]-Aspartate** | + | 0.20 | 0.85 | 139.06 | 92.25 | 20 | -12 | -13/-16 |
| **[$^{13}C_5$, $^{15}N_n$]-Glutamate** | + | 0.20 | 0.85 | 154.07 | 107.20 | 20 | -13 | -10/-19 |

| | | | | | | | | |
|---|---|---|---|---|---|---|---|---|
| **[$^{13}C_5$, $^{15}N_1$]-Proline** | + | 0.20 | 0.85 | 122.08 | 75.20 | 20 | -17 | -11/-20 |
| **[$^{13}C_3$, $^{15}N_1$]-Serine** | + | 0.20 | 0.85 | 110.06 | 63.20 | 20 | -14 | -10/-23 |
| **[$^{13}C_4$, $^{15}N_1$]-Threonine** | + | 0.20 | 0.85 | 125.08 | 78.20 | 20 | -12 | -12/-30 |
| **L-Proline** | + | 0.20 | 0.85 | 116.07 | 70.20 | 20 | -17 | -11/-20 |
| **L-Serine** | + | 0.20 | 0.85 | 106.05 | 60.20 | 20 | -14 | -10/-23 |
| **L-Threonine** | + | 0.20 | 0.85 | 120.07 | 74.20 | 20 | -12 | -12/-30 |
| **L-Asparagine** | + | 0.20 | 0.85 | 133.06 | 87.20 | 20 | -11 | -13/-16 |
| **L-Glutamine** | + | 0.20 | 0.85 | 147.08 | 130.15 | 20 | -15 | -10/-13 |
| **[$^{13}C_5$, $^{15}N_1$]-Valine** | + | 0.35 | 1.05 | 124.10 | 77.20 | 20 | -12 | -22/-28 |
| **L-Valine** | + | 0.35 | 1.05 | 118.09 | 72.20 | 20 | -12 | -22/-28 |
| **L-Isoleucine** | + | 0.85 | 1.70 | 132.10 | 86.15 | 122 | -11 | -20/-20 |
| **[$^{13}C_6$, $^{15}N_1$]-Isoleucine** | + | 0.85 | 1.70 | 139.12 | 92.25 | 122 | -11 | -20/-20 |
| **L-Leucine** | + | 0.85 | 1.70 | 132.10 | 86.15 | 122 | -11 | -20/-20 |
| **[$^{13}C_6$, $^{15}N_1$]-Leucine** | + | 0.85 | 1.70 | 139.12 | 92.25 | 122 | -11 | -20/-20 |
| **[$^{13}C_9$, $^{15}N_1$]-Phenylalanine** | + | 2.00 | 3.00 | 176.11 | 129.25 | 247 | -15 | -20/-20 |
| **L-Phenylalanine** | + | 2.00 | 3.00 | 166.09 | 120.20 | 247 | -15 | -20/-20 |
| **Cyanidin-3-O-glycoside** | + | 3.00 | 3.70 | 449.10 | 137.10 | 32 | -54 | -20/-14 |
| **Chlorogenic acid** | + | 3.00 | 3.70 | 355.10 | 163.15 | 32 | -15 | -23/-17 |
| **L-Tryptophan** | + | 3.00 | 3.70 | 205.10 | 188.10 | 32 | -10 | -20/-20 |
| **Apigenin-7-O-glycoside** | + | 3.40 | 4.10 | 431.10 | 268.10 | 32 | -35 | -21/-17 |
| **Apigenin-8-C-glycoside** | + | 3.40 | 4.10 | 431.10 | 311.20 | 32 | -24 | -30/-21 |
| **Luteolin-7-O-glycoside** | + | 3.40 | 4.10 | 447.09 | 285.20 | 32 | -26 | -22/-19 |
| **Luteolin-8-C-glycoside** | + | 3.40 | 4.10 | 447.09 | 327.15 | 32 | -24 | -30/-22 |

Abbreviations: CE - Collision energy

Table S3: Technical parameters used for quantification of the amino acids and secondary metabolites with Method S2 via LC-MS.

| Metabolite | Ionization mode | Start [min] | End [min] | Q1 [m/z] | Q3 [m/z] | Dwell time [ms] | CE | Q1/Q3 Pre Bias [V] |
|---|---|---|---|---|---|---|---|---|
| **Glycine** | + | 0.20 | 0.90 | 76.04 | 30.20 | 50 | -12 | -14/-11 |
| **[$^{13}C_2$, $^{15}N_1$]-Glycine** | + | 0.20 | 0.90 | 79.04 | 32.10 | 50 | -12 | -14/-11 |
| **L-Histidine** | + | 0.20 | 1.00 | 156.08 | 110.20 | 50 | -15 | -30/-21 |
| **[$^{13}C_6$, $^{15}N_3$]-Histidine** | + | 0.20 | 1.00 | 165.09 | 118.20 | 50 | -15 | -30/-21 |
| **[$^{13}C_5$, $^{15}N_1$]-Methionine** | + | 0.40 | 1.10 | 156.07 | 63.15 | 50 | -22 | -10/-24 |
| **L-Methionine** | + | 0.40 | 1.10 | 150.06 | 61.16 | 50 | -22 | -10/-24 |
| **Shikimic acid** | - | 0.40 | 1.10 | 173.05 | 93.10 | 23 | 15 | 12/21 |
| **[$^{13}C_9$, $^{15}N_1$]-Tyrosine** | + | 0.40 | 2.00 | 192.11 | 145.20 | 50 | -15 | -18/-14 |
| **L-Tyrosine** | + | 0.40 | 2.00 | 182.08 | 136.20 | 50 | -15 | -18/-14 |
| **Tyramine** | + | 0.40 | 2.00 | 138.09 | 121.15 | 23 | -15 | -20/-20 |
| **Apigenin** | - | 3.80 | 4.50 | 269.05 | 151.20 | 147 | 25 | 30/14 |
| **Luteolin** | - | 3.80 | 4.50 | 285.04 | 132.20 | 147 | 48 | 30/26 |

Abbreviations: CE - collision energy

Table S4: Technical parameters used for quantification of metabolites with Method S3 via LC-MS.

| Metabolite | Ionization mode | Start [min] | End [min] | Q1 [m/z] | Q3 [m/z] | Dwell time [ms] | CE | Q1/Q3 Pre Bias [V] |
|---|---|---|---|---|---|---|---|---|
| **SA** | - | 2.20 | 3.20 | 137.02 | 92.95 | 20 | 19 | 27/19 |
| **[$^2H_4$]-SA** | - | 2.20 | 3.20 | 141.05 | 97.00 | 20 | 19 | 27/15 |
| **ABA** | - | 2.20 | 3.20 | 263.13 | 153.10 | 20 | 13 | 18/29 |
| **[$^2H_6$]-ABA** | - | 2.20 | 3.20 | 269.17 | 159.20 | 20 | 13 | 18/30 |
| **JA** | - | 2.75 | 3.75 | 209.12 | 59.00 | 20 | 14 | 14/11 |
| **[$^2H_5$]-JA** | - | 2.75 | 3.75 | 214.15 | 62.00 | 20 | 14 | 14/10 |
| **JA-Ile** | - | 3.75 | 5.25 | 322.20 | 130.10 | 135 | 22 | 24/20 |

Abbreviations: ABA – Abscisic acid, CE - collision energy, JA – Jasmonic acid, JA-Ile – Jasmonic acid-isoleucine conjugate, SA – Salicylic acid

Table S5: Technical parameters used for quantification of the metabolites with Method S4 via LC-MS.

| Metabolite | Ionization mode | Start [min] | End [min] | Q1 [m/z] | Q3 [m/z] | Dwell time [ms] | CE | Q1/Q3 Pre Bias [V] |
|---|---|---|---|---|---|---|---|---|
| **Caffeic acid** | - | 1.95 | 2.70 | 179.03 | 135.05 | 20 | 18 | 12/26 |
| **Coumaric acid** | - | 2.30 | 3.05 | 163.04 | 119.10 | 20 | 17 | 11/11 |
| **Sinapic acid** | + | 2.45 | 3.15 | 225.08 | 207.15 | 20 | -9 | -15/-23 |
| **IAA** | + | 2.90 | 3.65 | 176.07 | 130.10 | 20 | -16 | -12/-23 |
| **[$^{13}C_6$]-IAA** | + | 2.90 | 3.65 | 182.09 | 136.25 | 20 | -16 | -12/-13 |

Abbreviations: CE - collision energy, IAA – Indole-3-acetic acid

Table S6: Technical parameters used for quantification of the metabolites with Method S5 via LC-MS.

| Metabolite | Ionization mode | Start [min] | End [min] | Q1 [m/z] | Q3 [m/z] | Dwell time [ms] | CE | Q1/Q3 Pre Bias [V] |
|---|---|---|---|---|---|---|---|---|
| **[$^2H_5$]-tZ** | + | 2.00 | 3.00 | 225.15 | 137.20 | 47 | -20 | -21/-22 |
| **tZ** | + | 2.00 | 3.50 | 220.12 | 136.10 | 47 | -20 | -20/-20 |
| **cZ** | + | 2.00 | 3.50 | 220.12 | 136.10 | 47 | -20 | -20/-20 |
| **tZR** | + | 3.75 | 5.25 | 352.16 | 220.15 | 30 | -20 | -20/-20 |
| **cZR** | + | 3.75 | 5.25 | 352.16 | 220.15 | 30 | -20 | -20/-20 |
| **iP** | + | 5.00 | 6.00 | 204.12 | 136.10 | 30 | -15 | -20/-20 |
| **[$^2H_6$]-iP** | + | 5.00 | 6.00 | 210.16 | 137.20 | 30 | -15 | -20/-20 |
| **iPR** | + | 6.00 | 7.00 | 336.17 | 204.20 | 47 | -20 | -20/-20 |
| **[$^2H_6$]-iPR** | + | 6.00 | 7.00 | 342.20 | 210.25 | 47 | -20 | -20/-20 |

Abbreviations: CE – Collision energy, cZ – cis-Zeatin, cZR – cis-Zeatin riboside, iP – Isopentenyladenine, iPR -Isopentenyladenine riboside, tZ – trans-Zeatin, tZR – trans-Zeatin riboside

Table S7: Primers used for RT-qPCR analysis of *SpUBP7* and reference gene (*SpaEF* and *SpGAPDH*) expression. The primers for the reference genes were used in previous publications (Wang et al. 2024; Höfer et al. 2024b).

| Gene | Gene ID | Forward Primer (5'-3') | Reverse Primer (5'-3') | Product size | Efficiency |
|---|---|---|---|---|---|
| ***SpGAPDH*** | *SpGA2022_054082* | AGCATCCAAGAAGGT GAAGATCGGC | TTGTAGTCGGTCGTGA TGAAGGGG | 132 bp | 104.4% |
| ***SpaEF*** | *SpGA2022_005771* | TCGAAGCCGGCATTT CCAAGGACG | TCGCCTTCGAGTACTT GGGTGTCG | 129 bp | 99.8% |
| ***SpUBP7*** | *SpGA2022_056000* | GACGCAGATTGGTCA ACCTTAGGTG | CCGGAACTGAATGTA GACACTGGATTG | 217 bp | 101.7% |

Table S8: Variation of compounds belonging to amino acid metabolism in 137 *S. polyrhiza* genotypes.

| Metabolite | Mean [µmol/g DW] | SE [µmol/g DW] | Range [µmol/g DW] | | Fold change | H2.c |
|---|---|---|---|---|---|---|
| | | | Minimum | Maximum | | |
| Glycine | 1.650 | 0.087 | 0.488 | 6.193 | 12.691 | 0.701 |
| L-Serine | 15.775 | 0.638 | 5.034 | 52.559 | 10.441 | $5.706 \cdot 10^{-9}$ |
| L-Alanine | 10.283 | 0.301 | 3.746 | 20.019 | 5.344 | 0.026 |
| L-Valine | 2.298 | 0.094 | 1.003 | 9.915 | 9.885 | 0.572 |
| L-Leucine | 0.579 | 0.022 | 0.232 | 1.786 | 7.698 | 0.771 |
| L-Aspartate | 24.468 | 0.622 | 10.529 | 49.306 | 4.683 | $1.336 \cdot 10^{-10}$ |
| L-Asparagine | 114.414 | 4.459 | 29.514 | 293.244 | 9.936 | $2.220 \cdot 10^{-15}$ |
| L-Methionine | 0.446 | 0.016 | 0.118 | 1.177 | 9.975 | 0.027 |
| L-Threonine | 6.759 | 0.276 | 2.187 | 19.990 | 9.140 | 0.187 |
| L-Isoleucine | 0.616 | 0.020 | 0.245 | 1.664 | 6.792 | 0.350 |
| L-Glutamine | 57.034 | 3.280 | 10.845 | 283.849 | 26.173 | 0.348 |
| L-Glutamate | 25.204 | 0.726 | 11.234 | 60.675 | 5.401 | $5.477 \cdot 10^{-10}$ |
| L-Arginine | 5.727 | 0.401 | 0.293 | 32.078 | 109.481 | 0.161 |
| L-Proline | 1.089 | 0.036 | 0.407 | 2.505 | 6.155 | 0.087 |
| L-Phenylalanine | 0.729 | 0.024 | 0.281 | 1.675 | 5.961 | 0.315 |
| L-Tyrosine | 0.285 | 0.008 | 0.106 | 0.684 | 6.453 | 0.591 |
| L-Tryptophan | 0.623 | 0.044 | 0.104 | 3.190 | 30.673 | 0.372 |
| Tyramine | 0.076 | 0.007 | 0.013 | 0.623 | 47.923 | 0.810 |

| | | | | | | |
|---|---|---|---|---|---|---|
| Shikimic acid | 1.798 | 0.080 | 0.695 | 9.173 | 13.199 | 0.732 |

Abbreviations: SE- standard error, H2.c – broad sense heritability according to Cullis et al. (2006)

Table S9: Variation of phytohormone concentrations in 137 *S. polyrhiza* genotypes.

| Metabolite | Mean [nmol/g DW] | SE [nmol/g DW] | Range [nmol/gDW] Minimum | Maximum | Fold change | H2.c |
|---|---|---|---|---|---|---|
| SA | 3.585 | 0.098 | 1.295 | 7.424 | 5.733 | 0.373 |
| ABA | 0.037 | 0.002 | 0.014 | 0.102 | 7.286 | $2.327 \cdot 10^{-7}$ |
| JA | 4.147 | 0.180 | 0.547 | 12.726 | 23.265 | 0.414 |
| JA-Ile | 1.760 | 0.074 | 0.223 | 4.471 | 20.049 | 0.382 |
| iP | 4.731 | 0.317 | 2.847 | 31.200 | 10.959 | 0.999 |
| iPR | 5.121 | 0.624 | 1.012 | 48.089 | 47.519 | 0.709 |
| tZ | 26.715 | 1.310 | 11.813 | 102.724 | 8.696 | 0.473 |
| tZR | 31.930 | 1.573 | 6.209 | 148.603 | 23.933 | 0.349 |
| cZ | 5.268 | 0.165 | 1.212 | 12.868 | 10.617 | 0.333 |
| cZR | 86.971 | 3.509 | 4.016 | 305.175 | 75.990 | 0.424 |
| IAA | 46.315 | 0.683 | 28.333 | 79.422 | 2.803 | 0.180 |

Abbreviations: ABA – Abscisic acid, cZ – cis-Zeatin, cZR – cis-Zeatin riboside, IAA – Indoleacetic acid, iP – Isopentenyladenine, iPR – Isopentenyladenine riboside, JA – Jasmonic acid, JA-Ile – Jasmonic acid-isoleucine conjugate, SA – Salicylic acid, tZ – trans-Zeatin, tZR – trans-Zeatin riboside, SE- standard error, H2.c – broad sense heritability according to Cullis et al. (2006)

Table S10: Variation of secondary metabolites concentrations in 137 *S. polyrhiza* genotypes.

| Metabolite | Mean [nmol/g DW] | SE [nmol/g DW] | Range [nmol/g DW] Minimum | Maximum | Fold change | H2.c |
|---|---|---|---|---|---|---|
| Chlorogenic acid | 2147.696 | 230.439 | 75.289 | 16181.370 | 214.923 | 0.979 |
| Coumaric acid | 10.461 | 0.555 | 5.078 | 49.489 | 9.746 | 0.529 |
| Caffeic acid | 3.784 | 0.485 | 1.029 | 60.979 | 59.260 | 0.927 |

| | | | | | | |
|---|---|---|---|---|---|---|
| Sinapic acid | 1.318 | 0.128 | 0.044 | 7.361 | 167.295 | 0.839 |
| Apigenin | 2.361 | 0.473 | 0.420 | 35.256 | 93.943 | 0.045 |
| Luteolin | 42.592 | 4.427 | 2.666 | 264.595 | 98.498 | 0.440 |
| Cyanidin-3-O-glycoside | 3972.074 | 273.743 | 403.464 | 17431.470 | 43.205 | 0.593 |
| Luteolin-8-C-glycoside | 15861.540 | 539.504 | 649.239 | 37498.920 | 57.758 | 0.112 |
| Luteolin-7-O-glycoside | 23596.070 | 780.300 | 256.907 | 47213.030 | 183.775 | 0.057 |
| Apigenin-8-C-glycoside | 11698.560 | 471.242 | 3696.448 | 32069.280 | 8.676 | 0.015 |
| Apigenin-7-O-glycoside | 5348.446 | 222.837 | 110.307 | 15238.690 | 138.148 | $1.294 \cdot 10^{-10}$ |

Abbreviations: SE- standard error, H2.c – broad sense heritability according to Cullis et al. (2006)

Table S11: Number of SVs and SNPs significantly associated with the corresponding parameters (Bonferroni-corrected *P*-Wald < 0.05). Parameters with no significant associations were not listed.

| **Parameter** | **Number significant markers-SVs** | **Number significant markers-SNPs** |
|---|---|---|
| RGR frond area | 1 | 0 |
| L-Arginine | 1 | 1 |
| L-Glutamine | 1 | 2 |
| L-Histidine | 1 | 3 |
| L-Leucine | 0 | 1 |
| L-Serine | 1 | 0 |
| L-Threonine | 0 | 1 |
| L-Tryptophan | 0 | 3 |
| L-Tyrosine | 0 | 6 |
| L-Valine | 0 | 2 |
| Shikimic acid | 0 | 7 |
| Tyramine | 0 | 3 |
| Chlorogenic acid | 1 | 5 |

| | | |
|---|---|---|
| Coumaric acid | 1 | 1 |
| Caffeic acid | 1 | 0 |
| Sinapic acid | 1 | 2 |
| Luteolin | 1 | 1 |
| Apigenin | 1 | 2 |
| Cyanidin-3-O-glycoside | 0 | 1 |
| iP | 3 | 12 |
| iPR | 0 | 7 |
| tZ | 1 | 6 |
| tZR | 0 | 6 |
| cZ | 1 | 0 |
| cZR | 0 | 1 |
| IAA | 0 | 2 |

Abbreviations: ABA – Abscisic acid, cZ – cis-Zeatin, cZR – cis-Zeatin riboside, IAA – Indoleacetic acid, iP – Isopentenyladenine, iPR – Isopentenyladenine riboside, JA – Jasmonic acid, JA-Ile – Jasmonic acid-isoleucine conjugate, SA – Salicylic acid, tZ – trans-Zeatin, tZR – trans-Zeatin riboside

Table S12: SVs significantly associated with metabolome and fitness data. The significance threshold was set to $4.23 \cdot 10^{-5}$ (Bonferroni corrected *P*-Wald value of 0.05 for 1182 markers).

| **Metabolite** | **Chromosome** | **Position** | **Associated gene locus and description** | **Location** | **Type of SV** | **Size of SV** | ***P*-Wald** |
|---|---|---|---|---|---|---|---|
| **L-Arginine** | 16 | 3514401 | - | Intergenic region: > 30 kbp away from next coding sequence | insertion | 56 bp | $1.32 \cdot 10^{-6}$ |
| **L-Glutamine, L-Serine** | 19 | 51003 | *SpGA2022_056000*: *SpUBP7* (Ubiquitin carboxyl-terminal hydrolase 7) | Intronic region (7th intron) of *SpUBP7* | deletion | 94 bp | $1.48 \cdot 10^{-5}$, $1.75 \cdot 10^{-5}$ |
| **L-Histidine** | 1 | 2805097 | *SpGA2022_002723* | 1814 bp upstream of SpGA2022_002723 | insertion | 97 bp | $6.44 \cdot 10^{-8}$ |

| | | | | | | | |
|---|---|---|---|---|---|---|---|
| **L-Leucine** | 14 | 4059780 | *SpGA2022_017710*: *SpROC1* (Homeobox-leucine zipper protein R) | 448 bp upstream of SpROC1 | deletion | 75 bp | $2.96 \cdot 10^{-5}$ |
| **Chlorogenic acid** | 10 | 421049 | *SpGA2022_013652* | 12392 bp upstream of *SpGA2022_013652* | deletion | 54 bp | $7.33 \cdot 10^{-7}$ |
| **Caffeic acid** | 12 | 4246070 | SpGA2022_016032: similar to At5g10620 (Putative RNA methyltransferase) | Intronic region (6th intron) of SpGA2022_016032 | duplication | 265 bp | $4.04 \cdot 10^{-5}$ |
| **Sinapic acid, Apigenin, Luteolin** | 16 | 4471839 | - | Intergenic region: > 30 kbp away from next coding sequence | deletion | 266 bp | $1.29 \cdot 10^{-9}$, $4.59 \cdot 10^{-7}$, $1.31 \cdot 10^{-6}$ |
| **iP** | 6 | 6329355 | *SpGA2022_009939*: Similar to Protein yippee-like | 10473 bp upstream of *SpGA2022_009939* | deletion | 54 bp | $2.16 \cdot 10^{-6}$ |
| **iP** | 13 | 2006566 | *SpGA2022_054972*: *SpNAP1* | Intronic region (5th intron) of *SpNAP1* | deletion | 74 bp | $2.98 \cdot 10^{-5}$ |
| **iP, tZ** | 18 | 95646 | *SpGA2022_019361*: *SpTSC10A* (3-dehydrosphinganine reductase) | 12371 bp upstream of *SpTSC10A* | deletion | 52 bp | $2.48 \cdot 10^{-8}$, $3.27 \cdot 10^{-5}$ |
| **cZ** | 13 | 1506652 | *SpGA2022_054953*: *SpSCAMP3* (Secretory carrier-associated membrane protein 3 | 3' UTR of *SpSCAMP3* | duplication | 76 bp | $6.73 \cdot 10^{-7}$ |
| **RGR frond area** | 6 | 4804486 | - | Intergenic region: > 30 kbp away from next coding sequence | insertion | 56 bp | $9.11 \cdot 10^{-6}$ |

Abbreviations: ABA – Abscisic acid, cZ – cis-Zeatin, cZR – cis-Zeatin riboside, IAA – Indoleacetic acid, iP – Isopentenyladenine, iPR – Isopentenyladenine riboside, JA – Jasmonic acid, JA-Ile – Jasmonic acid-isoleucine conjugate, SA – Salicylic acid, tZ – trans-Zeatin, tZR – trans-Zeatin riboside

Table S13: SNPs significantly associated with metabolome traits. The significance threshold was set to $1.14 \cdot 10^{-6}$ (Bonferroni corrected *P*-Wald value of 0.05 for 10057 markers).

| Parameter | Chromosome | Position | Alleles | Associated gene locus | Function | Location | *P*-Wald |
|---|---|---|---|---|---|---|---|
| **L-Arginine** | 12 | 1022717 | C/T | *SpGA2022_015481*: *SpNAT7* | Nucleobase ascorbate transporter 7 | Intronic region | $2.44 \cdot 10^{-8}$ |
| **L-Glutamine** | 2 | 5658111 | G/A | *SpGA2022_005065* | Unknown function | 2691 bp upstream | $4.08 \cdot 10^{-7}$ |
| **L-Glutamine** **L-Valine** | 4 | 6512246 | C/G | *SpGA2022_007596: SpLHCB5* | Chlorophyll a-b binding protein CP26, chloroplastic | Exonic region | $1.63 \cdot 10^{-7}$ $8.09 \cdot 10^{-7}$ |
| **L-Histidine** | 9 | 2006410 | G/A | *SpGA2022_053849*: *SpCYP71AU50* | Homolog of At1g51810 (Probable receptor-like protein kinase) | 7570 bp upstream | $2.15 \cdot 10^{-9}$ |
| **L-Histidine** | 8 | 2025860 | C/G | *SpGA2022_053495*: *SpPFK3* | ATP-dependent 6-phosphofructokinase 3 | 5334 bp upstream | $3.39 \cdot 10^{-9}$ |
| **L-Histidine,** **L-Tyrosine** | 20 | 3499796 | G/A | *SpGA2022_056315*: *SpPUB4* | U-box domain-containing protein 4 | 3404 bp upstream | $6.52 \cdot 10^{-8}$, $1.86 \cdot 10^{-8}$ |
| **L-Tryptophan** | 3 | 7646125 | T/A | *SpGA2022_006560* | Unknown function | Exonic region | $7.41 \cdot 10^{-7}$ |
| **L-Tryptophan** **L-Tyrosine** **L-Tyramine** | 14 | 3558175 | G/T | *SpGA2022_017644* | Unknown function | Intronic region | $7.41 \cdot 10^{-7}$ $3.52 \cdot 10^{-6}$ $1.39 \cdot 10^{-6}$ |
| **L-Tryptophan** **L-Tyrosine** **L-Tyramine** | 4 | 6512411 | G/A | *SpGA2022_007596*: *SpLHCB5* | Chloroplastic Chlorophyll a-b binding protein CP26 | Exonic region | $9.76 \cdot 10^{-7}$ |
| **L-Tyrosine** | 9 | 7058554 | A/G | *SpGA2022_054061*: *SpMYOB7* | Myosin binding protein 7 | 8600 bp upstream | $6.33 \cdot 10^{-7}$ |
| **L-Tyrosine** | 5 | 6307147 | G/A | *SpGA2022_008803*: *SpUGT89B1* | Flavonol 2-O-glycosyltransferase | 13387 bp upstream | $9.23 \cdot 10^{-7}$ |
| **L-Tyrosine** | 3 | 746125 | T/A | *SpGA2022_006560* | Unknown function | Exonic region | $3.52 \cdot 10^{-6}$ |
| **L-Valine** | 1 | 10088583 | G/A | *SpGA2022_003754* | Unknown function | Intronic region | $4.84 \cdot 10^{-8}$ |
| **Shikimic acid,** **iPR,** **tZR** | 5 | 1506810 | C/G | *SpGA2022_008160*: *SpRTNLB3* | Reticulon-like protein B3 | Exonic region | $7.97 \cdot 10^{-12}$, $1.22 \cdot 10^{-14}$, $1.36 \cdot 10^{-13}$ |
| **Shikimic acid,** **iPR,** **tZR** | 2 | 6978275 | G/A | *SpGA2022_005286*: *SpAGL62* | Transcription factor AGAMOUS | 707 bp upstream | $7.97 \cdot 10^{-12}$, $1.22 \cdot 10^{-14}$, $1.36 \cdot 10^{-13}$ |
| **Shikimic acid, iP,** **iPR** **tZ,** **tZR** | 8 | 2931769 | C/T | *SpGA2022_011897*: *SpLOB* | LOB (Lateral Organ Boundaries) | 12 bp upstream | $1.05 \cdot 10^{-9}$, $5.88 \cdot 10^{-19}$, $2.19 \cdot 10^{-8}$, $8.86 \cdot 10^{-12}$, $4.62 \cdot 10^{-8}$ |
| **Shikimic acid** | 1 | 4629613 | A/C | *SpGA2022_003090*: *SpERF071* | Ethylene-responsive transcription factor | Exonic region | $3.77 \cdot 10^{-6}$ |
| **Shikimic acid,** **iPR,** **tZR** | 12 | 660635 | C/G | - | Intergenic region > 60 kbp away from next coding sequence | - | $7.97 \cdot 10^{-12}$, $1.22 \cdot 10^{-14}$, $1.36 \cdot 10^{-13}$ |

| | | | | | | | |
|---|---|---|---|---|---|---|---|
| **Shikimic acid** | 14 | 1136868 | A/C | *SpGA2022_0055209* : *SpPEX19-1* | Peroxisome biogenesis protein 19-1 | Exonic region | $3.45 \cdot 10^{-6}$ |
| **Shikimic acid, iP** | 17 | 3839011 | A/T | *SpGA2022_019301*: *SpPID* | Protein kinase PINOID | 4235 bp downstream | $2.14 \cdot 10^{-6}$, $1.32 \cdot 10^{-6}$ |
| **Chlorogenic acid** | 3 | 3707844 | A/G | *SpGA2022_005797* | Unknown function | Exonic region | $7.92 \cdot 10^{-7}$ |
| **Chlorogenic acid** | 4 | 3248185 | A/G | *SpGA2022_007066*: *SpNOD19* | Nodulin19 | Intronic region | $7.69 \cdot 10^{-10}$ |
| **Chlorogenic acid** | 7 | 2101352 | G/C | *SpGA2022_010583* | Unknown function | Intronic region | $3.51 \cdot 10^{-6}$ |
| **Chlorogenic acid** | 9 | 2006394 | C/G | *SpGA2022_053849*: *SpCYP71AU50* | Cytochrome P450 71 | 7584 bp upstream | $1.34 \cdot 10^{-8}$ |
| **Chlorogenic acid** | 11 | 3813073 | G/A | *SpGA2022_015013*: Homolog of *At1g56130* | Probable LRR receptor-like serine/threonine-protein kinase | Exonic region | $3.30 \cdot 10^{-6}$ |
| **Coumaric acid** | 17 | 2596990 | T/C | *SpGA2022_055775*: *SpHAK12* | Putative potassium transporter 1 | 3400 bp upstream | $6.71 \cdot 10^{-7}$ |
| **Sinapic acid, Luteolin** | 1 | 9102932 | G/A | *SpGA2022_050859*: *SpTMEM184A* | Transmembrane protein 184A | 5772 bp downstream | $3.43 \cdot 10^{-6}$, $8.65 \cdot 10^{-7}$ |
| **Sinapic acid** | 3 | 2333724 | C/G | - | Intergenic region > 60 kbp away from next coding sequence | - | $3.88 \cdot 10^{-6}$ |
| **Apigenin** | 2 | 5607569 | C/T | *SpGA2022_051344* | Unknown function | 125 bp upstream | $2.78 \cdot 10^{-6}$ |
| **Apigenin** | 2 | 7491771 | C/T | *SpGA2022_005323* | Unknown function | 342 bp upstream | $4.31 \cdot 10^{-7}$ |
| **Cyanidin-3-O-glycoside** | 6 | 1885388 | A/G | - | Intergenic region > 40 kbp away from next coding sequence | - | $2.85 \cdot 10^{-7}$ |
| **iP** | 13 | 1671306 | C/T | *SpGA2022_054960*: *SpPNM1* | Pentratricopeptide repeat-containing protein PNM1 | 10531 bp upstream | $2.63 \cdot 10^{-9}$ |
| **iP** | 6 | 2254891 | T/C | - | Intergenic region > 20 kbp away from next coding sequence | - | $6.50 \cdot 10^{-9}$ |
| **iP** | 18 | 4022067 | A/C | *SpGA2022_019833*: homolog of *AT1G59675* | F-box containing protein | Exonic region | $4.34 \cdot 10^{-6}$ |
| **iP** | 9 | 6998734 | A/G | *SpGA2022_013565*: *SpBACOVA_02659* | Beta-glucosidase BoGH3B | Exonic region | $9.33 \cdot 10^{-7}$ |
| **iP** | 2 | 8321040 | G/A | *SpGA2022_051428*: *SpTOM1* | Tobamovirus multiplication protein 1 | Intronic region | $8.15 \cdot 10^{-7}$ |
| **iP** | 4 | 4459285 | G/A | *SpGA2022_007215*: *SpSPPL2* | Signal peptide peptidase-like 2 | Intronic region | $1.04 \cdot 10^{-6}$ |
| **iP** | 4 | 7453011 | T/C | *SpGA2022_007797* | Unknown function | 1567 bp upstream | $1.45 \cdot 10^{-6}$ |
| **iP, iPR,** | 5 | 4676913 | C/T | *SpGA2022_008667*: *SpHPL* | Fatty acid hydroperoxide lyase | 13696 bp upstream | $2.29 \cdot 10^{-9}$, $2.65 \cdot 10^{-9}$, |

| | | | | | | | |
|---|---|---|---|---|---|---|---|
| **tZ,**<br>**tZR** | | | | | | | $7.34 \cdot 10^{-11}$,<br>$1.14 \cdot 10^{-7}$ |
| **iP,**<br>**iPR,**<br>**tZ** | 4 | 6589981 | A/C | *SpGA2022_052203*:<br>*SpFRO6* | Ferric reduction oxidase 6 | 3182 bp upstream | $1.02 \cdot 10^{-13}$,<br>$4.75 \cdot 10^{-6}$,<br>$7.01 \cdot 10^{-8}$ |
| **iP,**<br>**iPR,**<br>**tZ,**<br>**tZR** | 7 | 4346941 | G/A | *SpGA2022_010966* | Unknown function | Intronic region | $2.49 \cdot 10^{-13}$,<br>$1.45 \cdot 10^{-8}$,<br>$2.90 \cdot 10^{-11}$,<br>$8.57 \cdot 10^{-9}$ |
| **tZ** | 6 | 2254891 | T/C | - | Intergenic region > 20 kbp away from next coding sequence | - | $2.46 \cdot 10^{-6}$ |
| **tZ** | 17 | 2596990 | T/C | *SpGA2022_055775*:<br>*SpHAK12* | Putative potassium transporter 1 | 3400 bp upstream | $5.33 \cdot 10^{-7}$ |
| **cZR** | 9 | 2886292 | T/C | *SpGA2022_053891*:<br>*SpPCMP-H12* | Pentratricopeptide repeat-containing protein | 6532 bp downstream | $3.87 \cdot 10^{-8}$ |
| **IAA** | 1 | 4153168 | G/A | *SpGA2022_002976*:<br>*SpMYBC1* | Transcription factor MYBC1 | Exonic region | $4.96 \cdot 10^{-7}$ |
| **IAA** | 1 | 7513300 | G/A | *SpGA2022_050798*:<br>*SpDPE2* | 4-alpha-glucanotransferase DPE2 | Exonic region | $2.44 \cdot 10^{-6}$ |

Abbreviations: ABA – Abscisic acid, cZ – cis-Zeatin, cZR – cis-Zeatin riboside, IAA – Indoleacetic acid, iP – Isopentenyladenine, iPR – Isopentenyladenine riboside, JA – Jasmonic acid, JA-Ile – Jasmonic acid-isoleucine conjugate, SA – Salicylic acid, tZ – trans-Zeatin, tZR – trans-Zeatin riboside
