## supplemental methods for "Genome-Wide Association Study of Metabolic Traits in the Duckweed *Spirodela polyrhiza*"

### Method S1

For measuring high abundant amino acids and secondary metabolites we used an injection volume of 0.5 µL. For elution solvents A and B were applied in a gradient mode with 0 min/2% solvent B, 1.5 min/2% solvent B, 3.5 min/100% solvent B, 4.5 min/100% solvent B, 5.0 min/2% solvent B and 6.0 min/2% solvent B. The mass spectrometer operated in multiple reaction monitoring mode (MRM) with settings reported in Table S2. All metabolites except of L-Asparagine, L-Glutamine, L-Tryptophan, Chlorogenic acid, Cyanidine-3-O-glycoside, Apigenine-7-O-glycoside, Apigenine-8-C-glycoside, Luteoline-7-O-glycoside, Luteoline-8-C-glycoside, Apigenine and Luteoline were quantified relative to their corresponding internal standards (Lab-X, with X being the amino acid). L-Asparagine was quantified relative to Lab-Aspartate and L-Glutamine relative to Lab-Glutamate. The other metabolites listed above were quantified relative to Lab-Phenylalanine.

### Method S2

We applied the same settings listed under Method S1. Since the amino acids and secondary metabolites quantified with this method were less abundant, we increased the injection volume to 5 µL. The metabolites Glycine, L-Histidine, L-Methionine and L-Tyrosine were quantified relative to their corresponding internal standards. Shikimic acid, Tyramine, Apigenine and Luteoline were quantified relative to the L-Phenylalanine standard. The MRM settings for the quantification are listed in Table S3.

### Method S3

For measuring phytohormones we used an injection volume of 5 µL. For elution, we applied solvents A and B in a gradient mode: 0.0 min/10% solvent B, 0.5 min/10% solvent B, 1.0 min/55% solvent B, 4.5 min/65% solvent B, 5.5 min/100% solvent B, 6.5 min/100% solvent B, 7.0 min/10% solvent B, 8.0 min/10% solvent B at a constant flow rate of 0.5 mL/min. Mass spectrometry measurements were conducted in MRM with settings listed in Table S4. The metabolites Abscisic acid (ABA), Jasmonic acid (JA) and Salicylic acid (SA) were quantified relative to their corresponding deuterated standards [$^{2}H_{6}$]-ABA, [$^{2}H_{5}$]-JA and [$^{2}H_{4}$]-SA (Olchemim, Olomouc, Czech Republic). We quantified the Jasmonic acid – Isoleucine conjugate (JA-Ile) relative to the [$D_{5}$]-JA standard.

**Method S4**

For measuring the metabolites IAA (Indoleacetic acid), Coumaric acid, Caffeic acid and Sinapic acid, we used an injection volume of 3 µL. MRM settings used for the quantification are listed in Table S5. Metabolites were eluted in a gradient mode with 0.0 min/10 % solvent B, 0.5 min/10% solvent B, 1.0 min/39% solvent B, 3.5 min/41% solvent B, 3.6 min/50% solvent B, 8.0 min/60% solvent B, 8.5 min/100% solvent B, 9.5 min/100% solvent B, 10.0 min/10% solvent B, 11.0 min/10% solvent B. All metabolites were quantified relative to the [$^{13}C_{6}$]-IAA internal standard.

**Method S5**

For quantifying cytokinins, we used an injection volume of 5 µL and followed the MRM settings listed in Table S6. The following gradient mode was applied for the mobile phase: 0.0 min/5 % solvent B, 0.5 min/5% solvent B, 0.7 min/15% solvent B, 3.5 min/25% solvent B, 6.5 min/70% solvent B, 6.7 min/100% solvent B, 7.7 min/100% solvent B, 8.0 min/5% solvent B, 9.0 min/5% solvent B. The metabolites trans-Zeatin (tZ), trans-Zeatin riboside (tZR), cis-Zeatin (cZ) and cis-Zeatin riboside (cZR) were quantified relative to the internal standard [$^{2}H_{5}$]-tZ. $N^{6}$ – Isopentenyladenine (iP) and $N^{6}$ – Isopentenyladenine riboside (iPR) were quantified relative to the [$^{2}H_{6}$] – iP standard.
