## supplemental figures for "Genome-Wide Association Study of Metabolic Traits in the Duckweed *Spirodela polyrhiza*"

**Supplementary Figures**

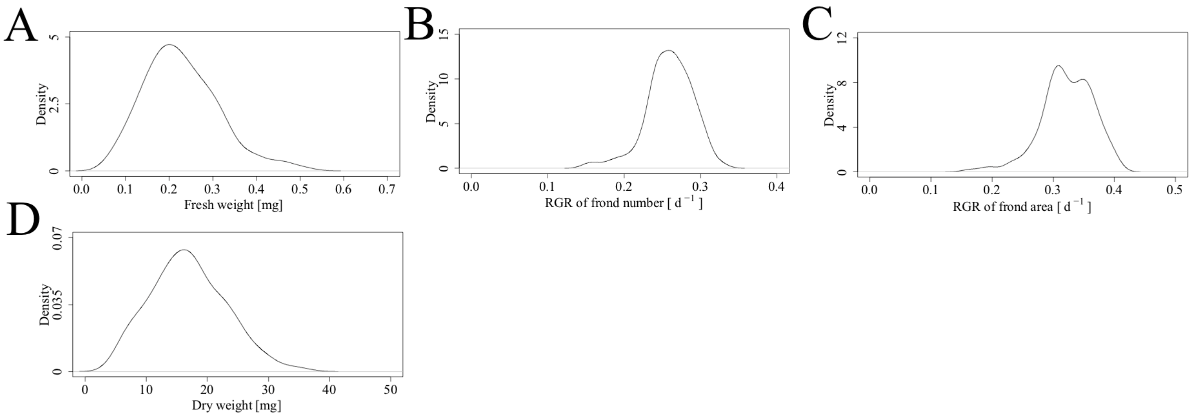


Figure S1: Kernel density distribution of the fitness parameters Fresh weight (A), RGR of frond number (B), RGR of frond area (C) and Dry weight (D) for 137 genotypes of *S. polyrhiza*.

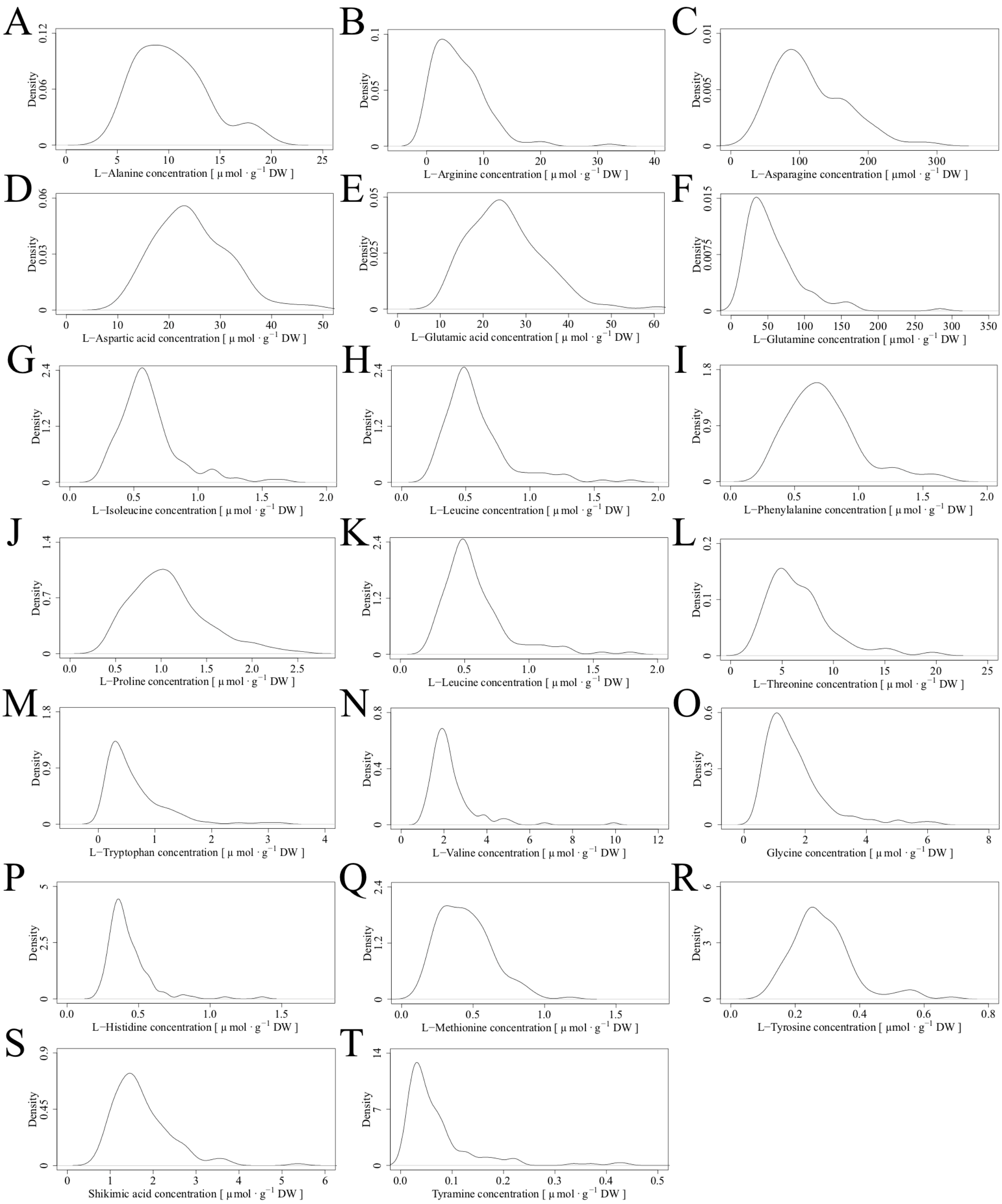


Figure S2: Kernel density distribution of amino acid concentrations measured for 137 genotypes of *S. polyrhiza*: L-Alanine (A), L-Arginine (B), L-Asparagine (C), L-Aspartic acid (D), L-Glutamic acid (E), L-Glutamine (F), L-Isoleucine (G), L-Leucine (H), L-Phenylalanine (I), L-Proline (J), L-Serine (K), L-Threonine (L), L-Tryptophan (M), L-Valine (N), Glycine (O), L-Histidine (P), L-Methionine (Q), L-Tyrosine (R), Shikimic acid (S) and Tyramine (T).

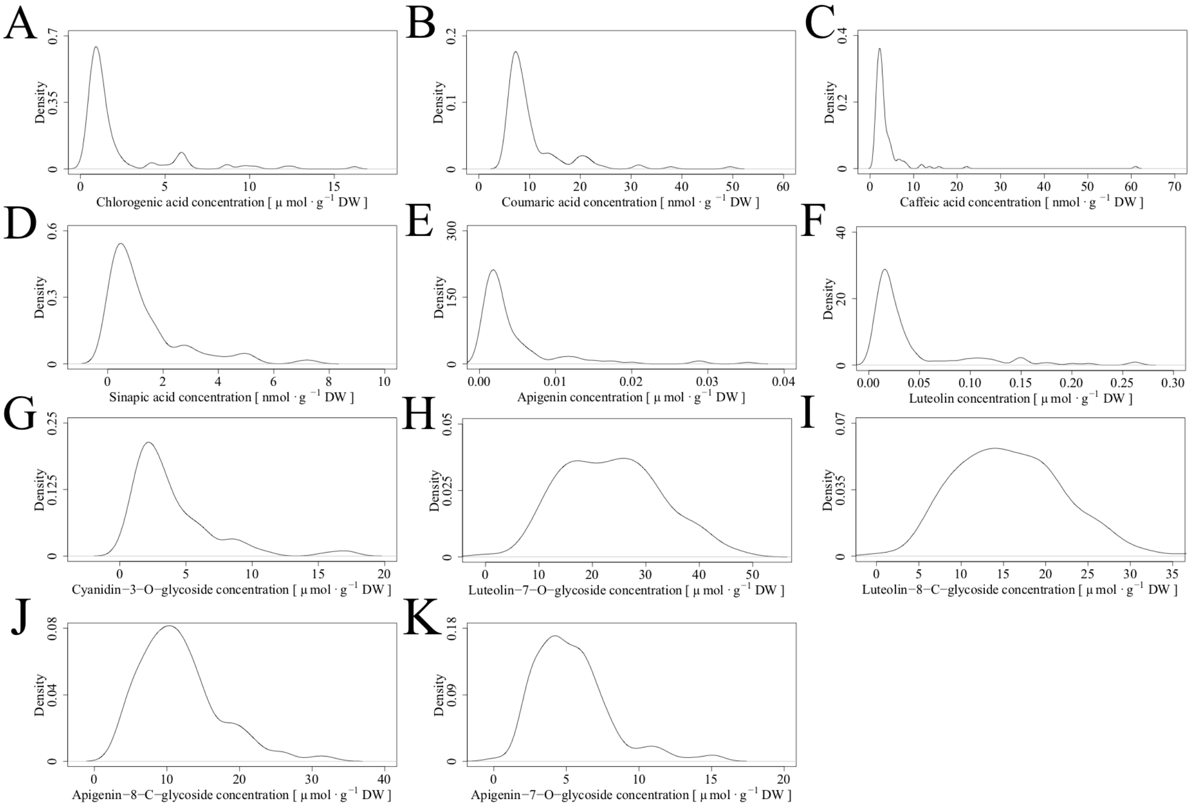


Figure S3: Kernel density distribution of secondary metabolite concentrations measured for 137 genotypes of *S. polyrhiza*: Chlorogenic acid (A), Coumaric acid (B), Caffeic acid (C), Sinapic acid (D), Apigenin (E), Luteolin (F), Cyanidin-3-O-glycoside (D), Luteolin-7-O-glycoside (E), Luteolin-8-C-glycoside (F), Apigenin-8-C-glycoside (G) and Apigenin-7-O-glycoside (H).

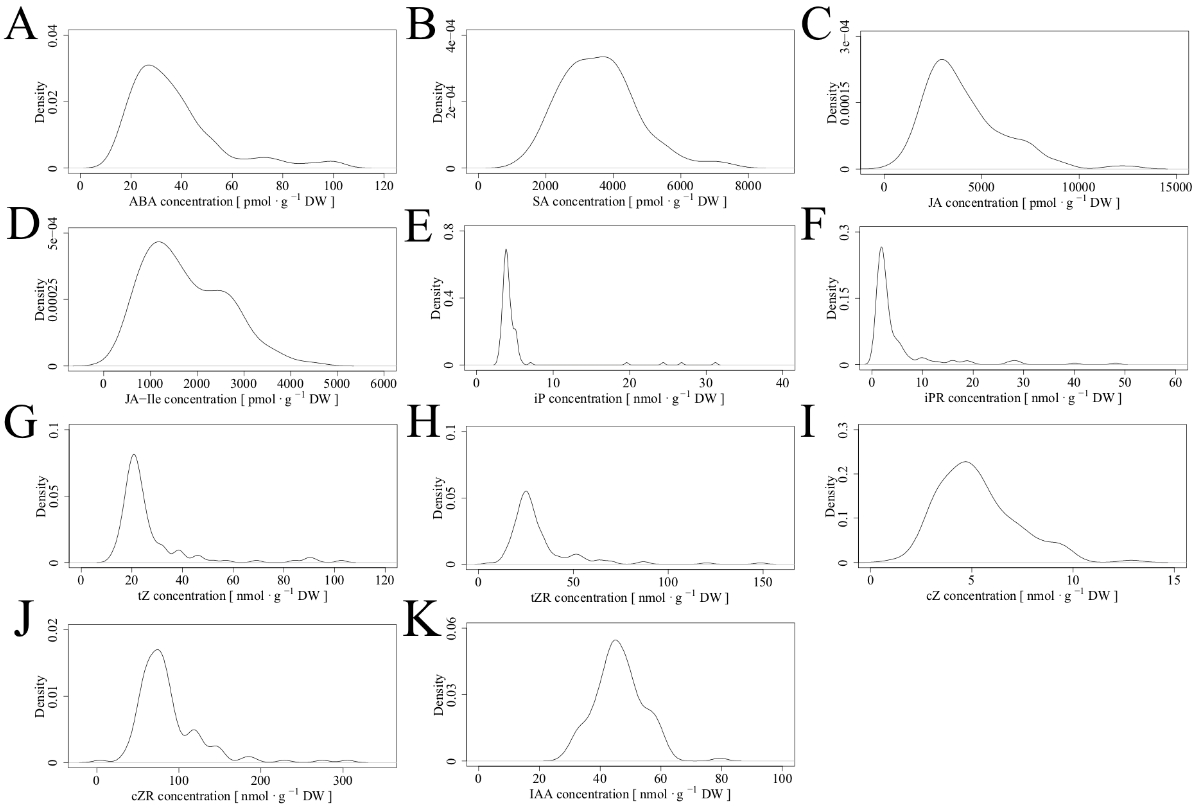


Figure S4: Kernel density distribution of phytohormone concentrations measured for 137 genotypes of *S. polyrhiza*: Abscisic acid (A), Salicylic acid (B), Jasmonic acid (C), Jasmonic acid-Isoleucine conjugate (D), $N^6$ – Isopentenyladenine (E), $N^6$ – Isopentenyladenine riboside (F), trans-Zeatin (G), trans-Zeatin riboside (H), cis-Zeatin (I), cis-Zeatin riboside (J), Indole-3-acetic acid (K).

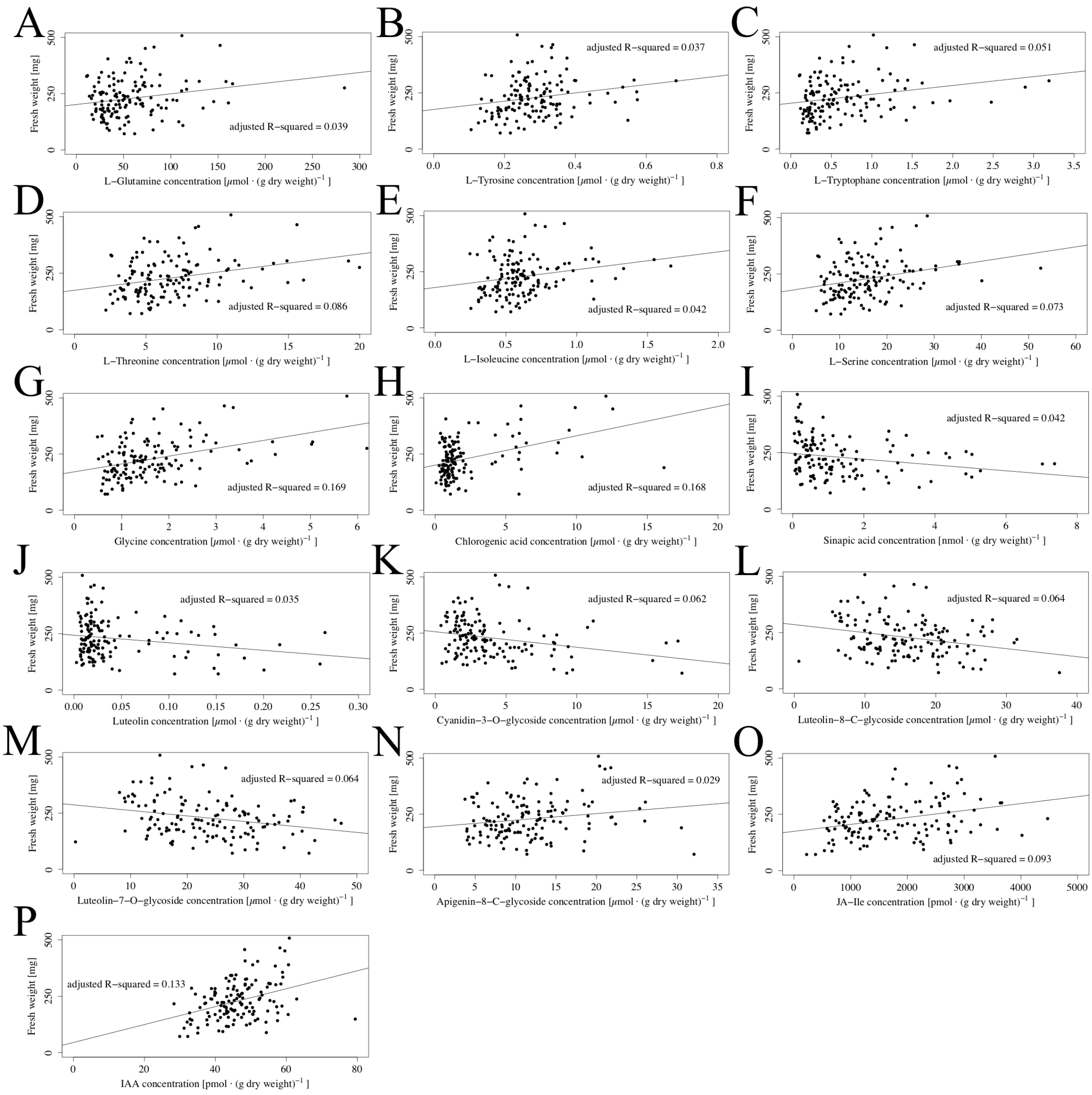


Figure S5: Scatterplots of all significant correlations (F-test, *P*-value ≤ 0.05) of fresh weight with individual free metabolite contents. Fresh weight significantly correlated with contents of L-Glutamine (A), L-Tyrosine (B), L-Tryptophane (C), L-Threonine (D), L-Isoleucine (E), L-Serine (F), Glycine (G), Chlorogenic acid (H), Sinapic acid (I), Luteolin (J), Cyanidin-3-O-glycoside (K), Luteolin-8-C-glycoside (L), Luteolin-7-O-glycoside (M), Apigenin-8-C-glycoside (N), Jasmonic acid – Isoleucine (O), Indole-3-acetic acid (P)

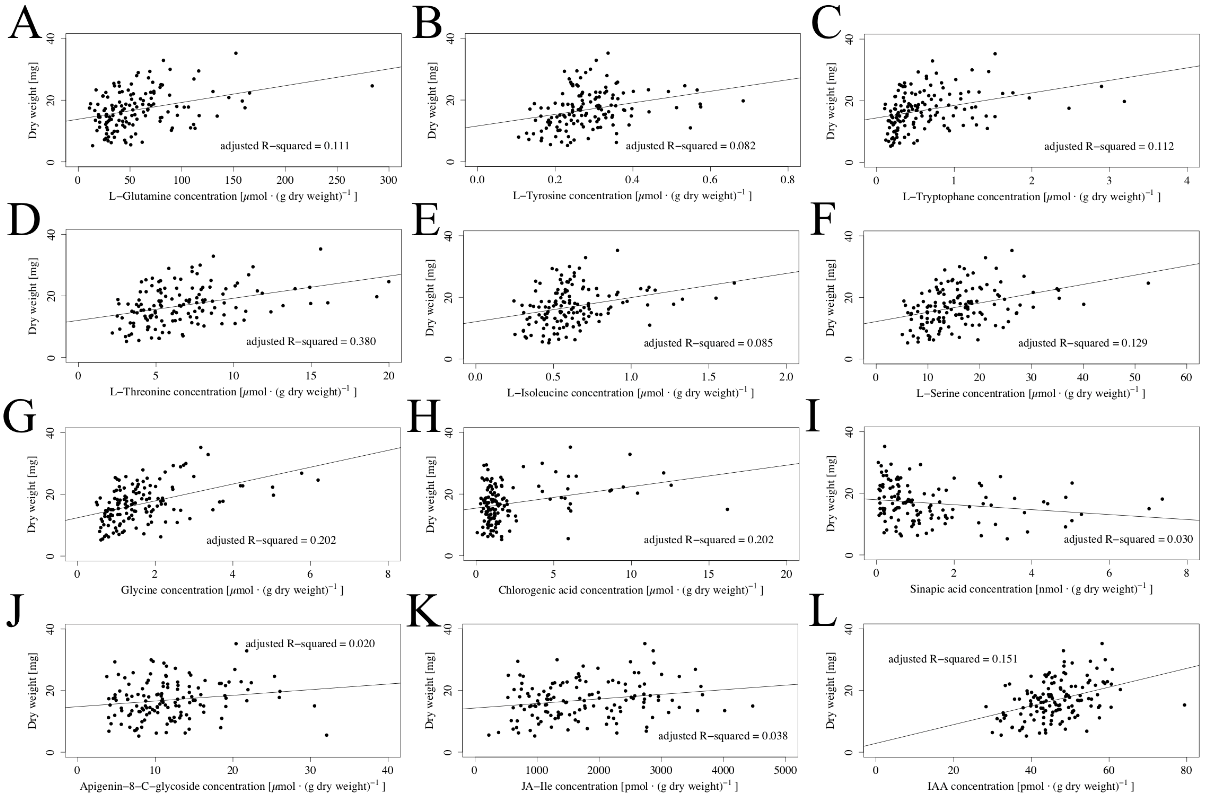


Figure S6: Scatterplots showing all identified significant correlations (F-test, *P*-value ≤ 0.05) of individual free metabolite contents with dry weight. Dry weight was significantly correlated with levels of L-Glutamine (A), L-Tyrosine (B), L-Tryptophane (C), L-Threonine (D), L-Isoleucine (E), L-Serine (F), Glycine (G), Chlorogenic acid (H), Sinapic acid (I), Apigenin-8-C-glycoside (J), Jasmonic acid – Isoleucine (K), Indole-3-acetic acid (L).

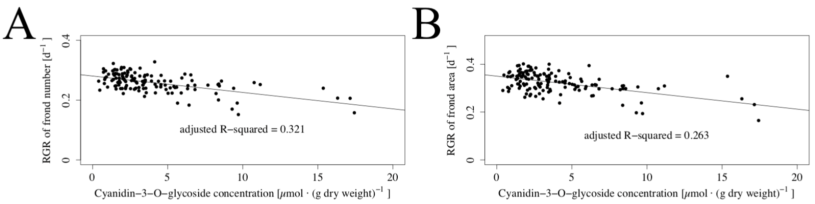


Figure S7: Correlation of growth with contents of Cyanidin-3-O-glycoside. Both metabolites showed strong negative correlation patterns with RGR of frond number (Pearson, ρ = -0.57) (A) and RGR of frond area (Pearson, ρ = -0.52) (B).

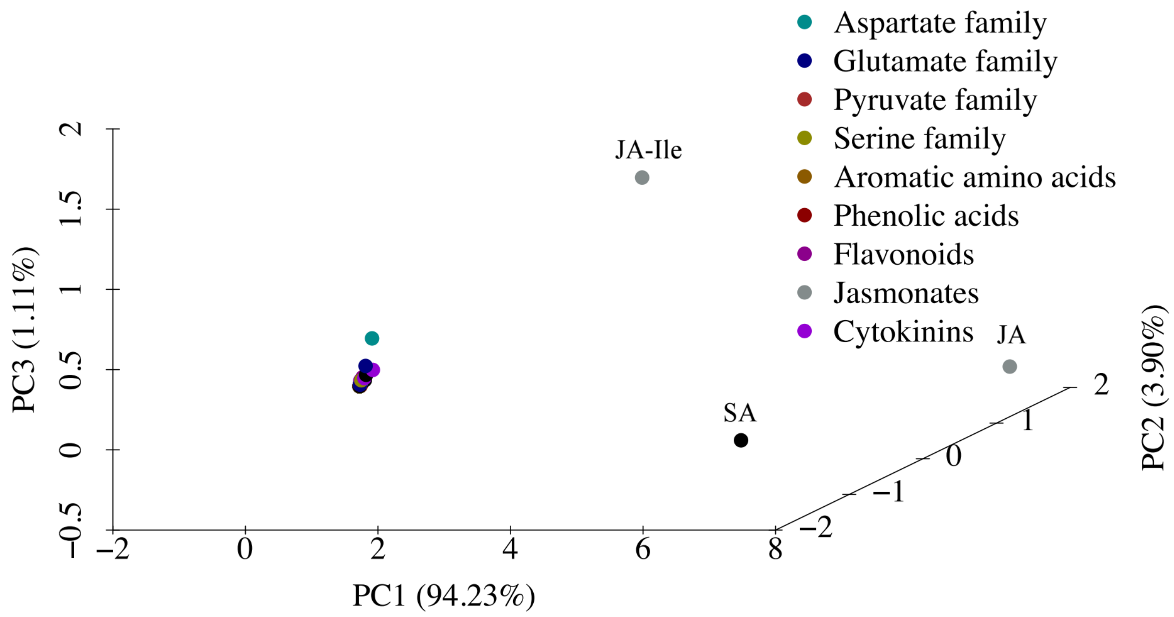


Figure S8: PCA on levels of 42 metabolites, that were categorized into nine groups. Contents of all metabolites except that of Salicylic acid (SA) and Jasmonates (JA and JA-Ile) formed a cluster.
